# Differential cell survival outcomes in response to diverse amino acid stress

**DOI:** 10.1101/2025.04.01.646536

**Authors:** Marion Russier, Alessandra Fiore, Annette Groß, Maria Tanzer, Assa Yeroslavitz, Matthias Mann, Peter J. Murray

## Abstract

Amino acid (AA) detection is fundamental for cellular function, balancing translation demands, biochemical pathways, and signaling networks. While the GCN2 and mTORC1 pathways are known to regulate AA sensing, the global cellular response to AA deprivation remains poorly understood, particularly in non-transformed cells, which may exhibit distinct adaptive strategies compared to cancer cells. Here we employed murine pluripotent embryonic stem (ES) cells as a model system to dissect responses to AA stress. Using multi-omics analyses over an extended time course, we examined the effects of arginine (Arg) and leucine (Leu) deprivation and uncovered a broad array of proteomic, phosphoproteomic, transcriptomic, and metabolomic adaptations, including an increase in lysosome production, all occurring without lethality. We found that Arg or Leu starvation induces reversible cell cycle exit, promoting a quiescent state that enhances resistance to cytotoxic stressors. By contrast, cysteine (Cys) and threonine (Thr) deprivation led to cell death via distinct pathways: ferroptosis for Cys starvation, while Thr deprivation triggered a previously uncharacterized form of cell death, which could be entirely suppressed by methionine (Met) co-starvation, mTOR or translational inhibition. These findings suggest that ES cells implement specialized survival strategies in response to different AA limitations, highlighting their ability to reprogram cellular biochemistry under nutrient stress.

## Introduction

Cells continuously monitor amino acid (AA) availability to regulate metabolism, translation, and growth^1–3^. AAs are sourced from different “pools” including external uptake via solute carrier (SLC) transporters, intracellular reservoirs such as lysosomes, biosynthetic pathways, and protein degradation. In transformed cells, increased proliferation alters AA demands, leading to adaptive strategies to obtain AA such as enhanced autophagy and ribophagy, increased solute transporter expression, uptake of abundant serum proteins ^4–9^ and catabolism of local matrix proteins or oligopeptides^10,11^ or secretion of specific AAs^12^. Key outstanding questions in AA metabolism include how cells detect individual AAs, assess their proportional balance, and integrate this information to regulate cellular pathways.

Two major AA-sensing pathways, GCN2 and mTORC1, mediate adaptive responses to AA availability^13–18^. GCN2 activation, triggered by uncharged tRNAs at stalled ribosomes, initiates the integrated stress response (ISR), reducing translation through eIF2α phosphorylation^19–23^. Conversely, mTORC1, activated by AA-binding proteins such as Sestrin-2 (Leu sensor) and Castor-2 (Arg sensor), promotes anabolic metabolism, translation, and ribosome biogenesis^13,15–18,24–28^. These pathways operate antagonistically, with GCN2 sensing AA scarcity and mTORC1 detecting AA abundance. Despite advances in understanding these pathways, major gaps remain in elucidating how mammalian cells collectively detect and prioritize AAs under physiological conditions^29–31^.

AA sensing and metabolic adaptation are particularly relevant in cancer, where specific AAs become metabolic dependencies. Moreover, AA limitation can drive cells into a quiescent state, rendering them vulnerable to metabolic perturbations^32–34^. However, AA sensing is highly context-dependent, with different cell types exhibiting unique responses^35^. Without a defined “baseline” model, interpreting cell-specific metabolic adaptations remains challenging. Here, we use murine embryonic stem (ES) cells as a non-transformed, rapidly proliferating model system (∼8-10 h division time) to investigate fundamental responses to AA limitation. As ES cells are pluripotent and non-malignant, they provide an ideal system to elucidate core mechanisms underlying cellular adaptation to AA stress, independent of oncogenic transformation.

Our study aimed to gain an understanding of how non-transformed cells sense and respond to AA deprivation in a global way by leveraging high-resolution proteomic, transcriptomic, and metabolomic approaches. By systematically comparing ES cell responses to deprivation of different AAs, we found both common and distinct mechanisms govern cellular adaptation to AA stress, including cell survival pathways.

## Results

### Global remodeling of cell state during long term AA starvation

To gain insights into how ES cells negotiate AA starvation, we employed a global, time-resolved multi-parametric approach, using deprivation of the essential AAs Arg or Leu, both of which activate mTORC1. Within hours of starvation, ES cells exited the cell cycle and showed a marked reduction in overall protein translation (**Fig. 1a, b**). Importantly, ES cells tolerated Arg or Leu deprivation stress reversibly; even after 48 h of Arg or Leu deprivation, starved ES cells remained viable (**Fig. S1a**). Re-addition of either Arg or Leu triggered immediate re-entry into the cell cycle as measured by live cell imaging (**Fig. 1c**). Using quantitative mass spectrometry over 1, 3, and 10 h of starvation revealed that, aside from Arg or Leu, the intracellular levels of other AAs remained largely stable—with the exception of proline, which increased in both conditions (**Fig. 1d**). The reasons behind this increase in proline remain unclear but may point to a yet to be understood role in AA stress adaptation^36^.

**Fig. 1.**
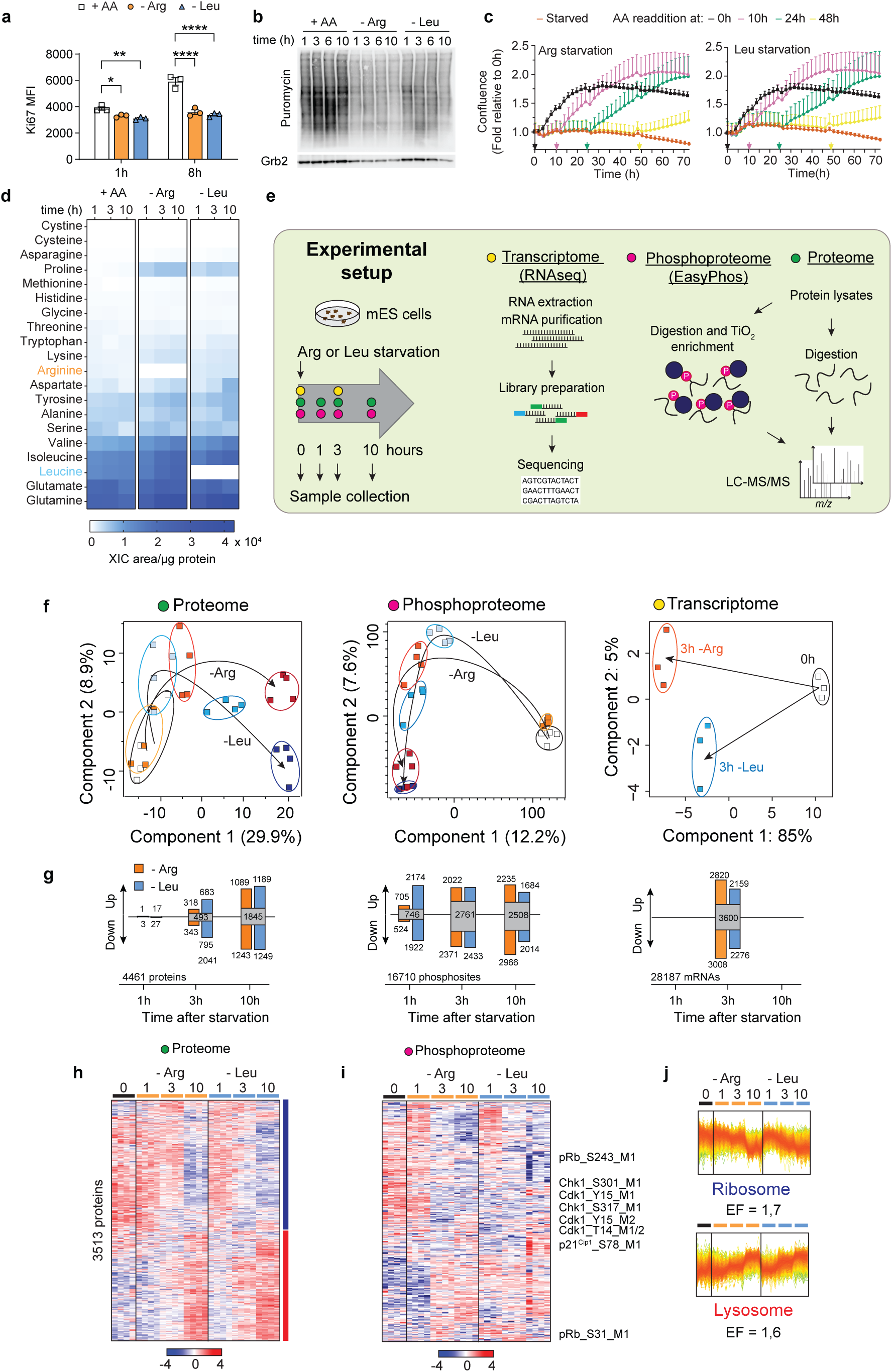
| Time-resolved multi-omics profiling of the AA starvation response. **a**, ES cells were starved of Arg or Leu and cell proliferation was assessed by Ki67 expression and intracellular flow cytometry. As a control, medium containing all AAs was used. Data is shown as the mean of fluorescence intensity (MFI) ± SEM, n=3. Statistical significance was determined by 2-way ANOVA followed by Dunnett’s multiple comparisons: * *p* < 0.05, ** *p* < 0.01, **** *p* < 0.0001. **b**, Newly synthesized proteins were labelled by puromycin incorporation and revealed by immunoblotting. **c**, Cell proliferation during Arg or Leu deprivation and re-supplementation at the indicated time points (arrows). Cell confluence was monitored by live imaging, normalized to 0 h and shown as mean ± SEM, n=3. **d**, Heatmap of all amino acids detected by LC-MS-based metabolomics in the cell lysates at 1, 3 and 10 h after Arg or Leu starvation. The data was calculated as the mean of the extracted ion chromatogram (XIC) per area of 3 replicates normalized to the total amount of protein. **e**, Schematic of the experimental setup and workflow of the omics data analysis. **f**, Principal component analysis (PCA) assessing temporal dynamics during Arg (orange, red shades) and Leu (blue shades) starvation. Each square represents a sample collected at a particular time point, with lighter and darker shades denoting early and late time points, respectively. **g**, Summary statistics of proteins, phosphosites and transcripts (read counts > 1) significantly up-or down-regulated at the indicated time points during Arg and Leu starvation, as determined by multiple *t*-tests relative to 0 h (proteome, phosphoproteome, and FDR < 5%; transcriptome: Wald test, Benjamini Hochberg-adjusted *p* value (FDR) < 5%). **h-i**, Heatmap of z-scored intensities of the proteome (h) and phosphoproteome (i). **h**, The proteome was analyzed by hierarchical cluster analysis (Euclidean distance) of all significantly regulated proteins (3513) compared to all identified (4461) (one-way ANOVA, *s0* = 0.1, FDR < 5%). Profiles of each cluster based on protein temporal dynamics are color-coded according to their distance from the respective cluster center. **i**, The phosphoproteome was analyzed by hierarchical cluster analysis (Euclidean distance) and represents all significantly regulated sites (8715) (one-way ANOVA, *s0* = 0.1, FDR < 5%). **j**, Fisher’s exact test on protein clusters with ribosome and lysosome KEGG annotations (background is all identified proteins in the dataset, cut-off *p* < 0.002). The enrichment factors (EF) are displayed. Top (ribosome) and bottom (lysosome) panels corresponding to blue and red clusters in **h**, respectively.

To gain a global view of how ES cells adapted their proteomes to AA stress, we deprived ES cells of Arg or Leu for 1, 3, or 10 h and profiled protein and phosphoprotein abundance in parallel with RNAseq analysis at 3 h, when cells ceased proliferation (**Fig. 1e**). Principal component analysis (PCA) revealed a broad reorganization of the proteome, phosphoproteome, and transcriptome during AA starvation (**Fig. 1f**). Changes in the proteome were gradual across the time course reaching a maximum of >3000 proteins significantly regulated by 10 h of starvation (**Fig. 1g**). Using hierarchical clustering, we identified two main temporal clusters: ∼ half the proteins decreased overtime (enriched with ribosomal components), while another group of proteins increased, particularly proteins associated with lysosome biogenesis and function (**Fig. 1h-j**). This pattern reflected a reduced need for ribosomal activity and an increase reliance on lysosome function, consistent with other results using murine ES cells^37^. Within the phosphoproteome, we detected thousands of phosphorylation events (**Fig. 1i**). Many of these modifications were linked to cell cycle control, particularly involving cyclin-dependent kinases (CDK), which is consistent with the shutdown of proliferation in cells existing the cell cycle. At the mRNA level, we observed 8-10,000 transcripts were up- and down-regulated relative to unstarved cells (**Fig. 1g**, right panel). Given the overall reduction of translation during AA starvation (**Fig. 1b**), the transcriptomes and proteomes were largely uncorrelated (**Fig. S2a, b**) This likely reflects a shift from bulk protein synthesis to the selective translation of stress-protective factors and the stabilization of specific proteins, alongside the activation of specialized transcriptional programs. Together, these findings illustrate that the adaptative response to long term nutrient stress of single essential AAs is a multi-level process involving extensive transcriptional, translational and post-translational reprogramming to achieve a temporary quiescent state.

GCN2 (encoded by *Eif2ak4*) is activated by limiting AA amounts. To understand the contribution of GCN2 to the broad re-wiring of the transcriptome in Leu or Arg starved ES cells, we generated *Eif2ak4*-deficient ES cells by Crispr-Cas9 targeting (**Fig. S1b**). As expected, activation of phosphorylation of GCN2 at the key T889 site was induced by deprivation of Arg or Leu, followed by the activation of the ISR as measured by ATF4 expression (**Fig. S1c**). We used the *Eif2ak4*-deficient ES to perform RNAseq in comparison to control ES cells. We observed that GCN2 (via its downstream mediators such as ATF4) controlled a substantial fraction of transcripts regulated by AA starvation, such as Txnip1 and Ddit3 (also known as CHOP) many of which are known targets of the integrated stress response (ISR) controlled by GCN2^14,38,39^ (**Fig. S1d, e**). However, many classes of regulated transcripts were independent of GCN2, including lysosome-associated mRNAs and many transcripts encoding metabolic enzymes (**Fig. S1d**). Therefore, starved ES cells re-wired their transcriptional program only part of which is GCN2-dependent, suggesting other pathways contribute to the overall AA stress response.

Although Arg and Leu are sensed by different upstream regulators—such as Sestrin-2 for Leu and Castor-2 for Arg—and are known to differentially affect ribosome pausing and the ISR^40^, our data from ES cells indicate a remarkably similar adaptive response to starvation of either AA. Quantitative comparisons of proteomic (10 hours) and transcriptomic (3 hours) data from Arg- and Leu-starved cells showed linear correlations (**Fig. S3a, b**), a trend that extended to the phosphoproteome across all time points (**Fig. S3c**) and to the specific proteins enriched during starvation (**Fig. S3d**). These findings suggest that in ES cells, Arg and Leu deprivation converge on a common adaptive program. This finding contrasts with observations in transformed cells where Arg deprivation more robustly induces ribosome pausing and GCN2 activation compared to Leu starvation^40^.

### Metabolic adaptation in the quiescent state

Given that the transcriptome and proteome results showed thousands of changes in response to Arg or Leu starvation, we next considered how ES cells globally adapted their metabolism once they entered a quiescent state. After an extended Arg or Leu starvation (17 h), the oxygen consumption rate remained near the assay baseline (**Fig. 2a**) and the mitochondrial membrane potential (reflecting mitochondrial activity) was indistinguishable from that of proliferating cells (**Fig. 2b**). Key metabolic intermediates (e.g. glucose-6-phosphate and succinate) were reduced (**Fig. 2c**), and this paralleled a decline in the NADH/NAD^+^ and NADPH/NADP^+^ ratios (**Fig. 2d**), while, the ATP/ADP ratio increased (**Fig. 2e**). In line with the transcriptomic and proteomic correlations, we did not observe any significant differences between Leu versus Arg starvation. These metabolic changes, together with preserved AA levels (**Fig. 1d**) and rapid cell cycle re-entry upon nutrient restoration (**Fig. 1c**), suggest that ES cells adopt a metabolic state that is poised for rapid recovery. Interestingly, ∼50% of key enzymes involved in central metabolism declined at the protein level. However, few corresponding mRNAs changes (only 3/36) were detected (**Fig. 2f, g**), reflecting the non-correlation between the AA starvation transcriptome-proteome (**Fig. S2**). When considered collectively, the metabolic status of AA starved cells was consistent with an overall decline in central metabolism, defined here by the pentose phosphate pathways or PPP, glycolysis, lactate production and the TCA cycle.

**Figure 2.**
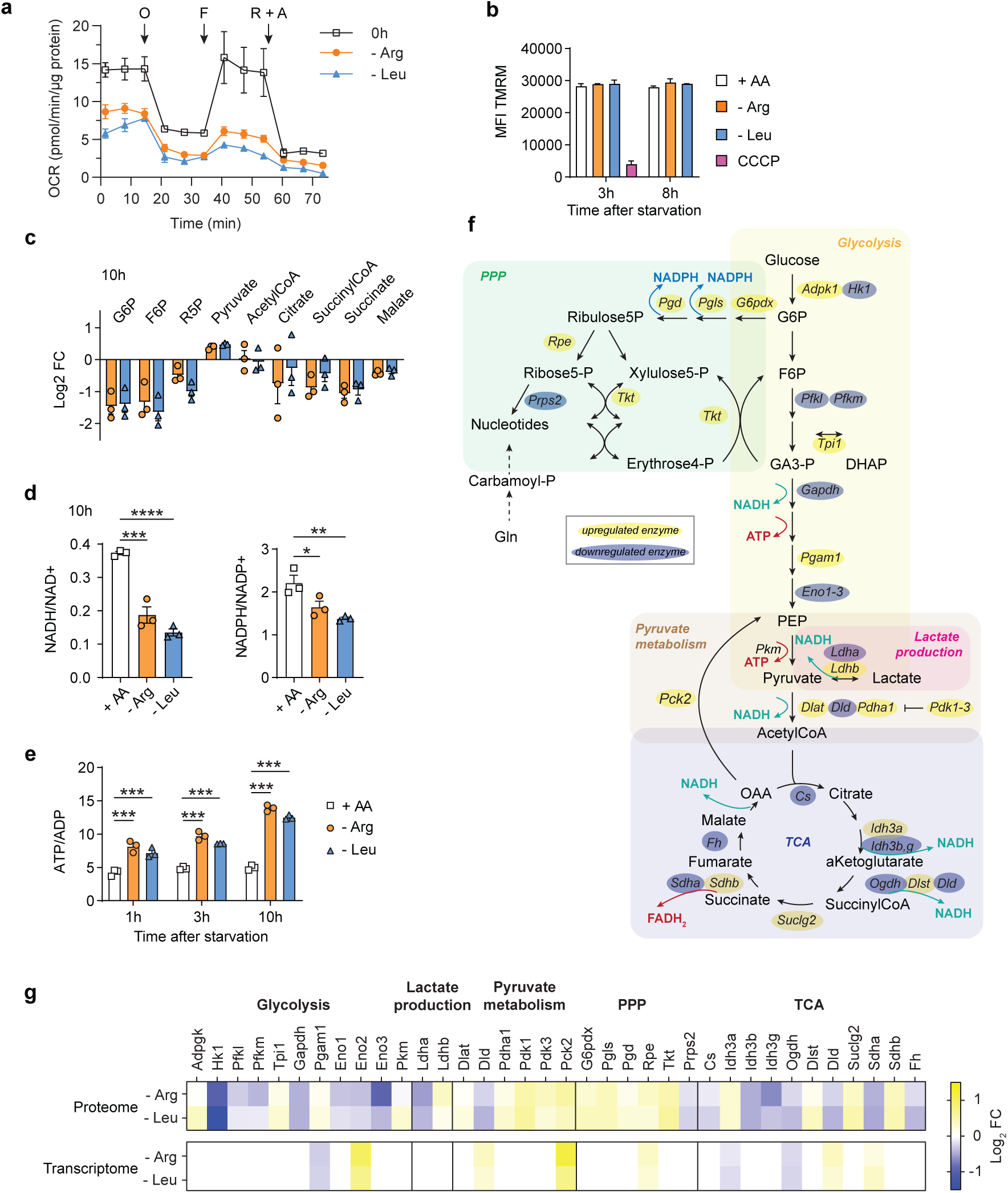
| Metabolic adaptation during the amino acid starvation response. **a**, Mitochondrial respiration of cells is reduced during prolonged amino acid deprivation. Cells were starved of Arg or Leu overnight (17 h) and assessed by Seahorse. Oxygen consumption rate (OCR) measurements were normalized to the amount of protein in the samples and shown as mean ± SEM of 3 replicates. O, oligomycin, F, FCCP, R+A, AntimycinA + rotenone. **b**, Mitochondrial potential is unaffected during Arg or Leu limitation, as assessed by TMRM fluorescent probe. As a control, 50 µM CCCP was added to inhibit oxidative phosphorylation. The results display the mean ± SEM of a representative experiment with 2 replicates. **c-e**, HPLC-MS-based metabolomics revealed a decline in central metabolism after Arg and Leu starvation. The changes in the abundance of the main metabolic intermediates 10h after starvation are displayed as mean of the fold change of Arg- or Leu-samples compared to the unstarved (+ AA) group (n = 3) (**c**). NADH/NAD^+^, NADPH/NADP^+^ (**d**) and ATP/ADP (**e**) ratios are shown as mean ± SEM of 3 replicates and calculated at 10h (**d**) or the indicated time points (**e**) after Arg or Leu starvation. Significant differences were obtained by one-way or two-way ANOVA followed by Dunnett’s multiple comparisons at each time point: * *p* < 0.05, ** *p*< 0.01, **** *p* < 0.0001. **f**, Scheme of the metabolic pathways summarizing the metabolic and related proteomic changes. The metabolic enzymes are highlighted according to the proteome dataset: in yellow or blue whether they are upregulated or downregulated after 10h of Arg or Leu starvation compared to 0h, respectively. **g**, Heatmap of the proteome and transcriptome changes (log_2_ fold change compared to the unstarved group) associated with metabolic adaptation.

### Cysteine or threonine deprivation causes distinct forms of cell death

We next considered if we could correlate the proteomic, transcriptomic and metabolic changes we quantified with Arg and Leu limitation to starvation to two AAs needed for ES functionality beyond translation: Cys and Thr. Cys is required for glutathione (GSH) biosynthesis, translation, trans-sulfuration, Fe-S biogenesis and other core redox-related cellular functions^41^. GSH is a core redox regulator essential for stress protection, including safeguarding against ferroptosis.^42^. Upon Cys removal, ES cells began to die after ∼10 hr (**Fig. 3a, b**). Death caused by the absence of Cys was rescued by the addition of free radical scavengers Ferrostatin-1 (Fer-1) and N-acetyl-cysteine (NAC) but not by inhibition of caspase-dependent apoptosis by z-VAD-fmk or necroptosis by Nec1s (**Fig. 3a, b**) indicating that Cys starvation induced ferroptosis in ES cells.

**Figure 3.**
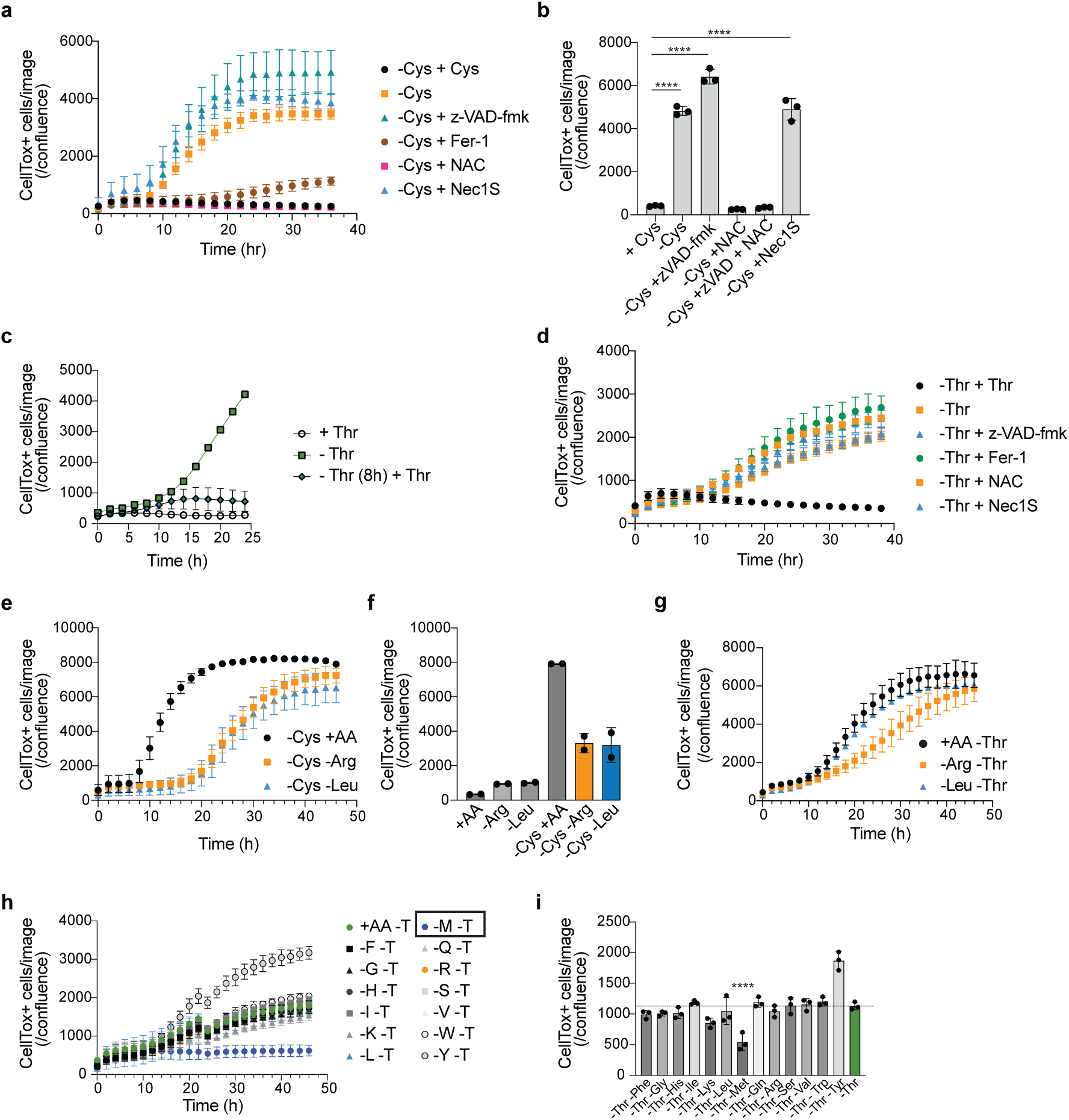
| ES cells are sensitive to Cys withdrawal. **a**, Quantification of cell death induced by Cys deprivation in the presence or absence of cell death inhibitors. Note that death induced by Cys starvation is blocked by re-addition of Cys, Fer-1 or NAC. A representative experiment with 2 technical replicates is shown. The data was calculated as a ratio of CellTox^+^ green object counts per image to the cell confluence area. **b**, Summary quantification of cell death induced by Cys deprivation in the presence or absence of cell death inhibitors at 24 h for each condition. **** *p* < 0.0001 Dunnett’s multiple comparison test after ordinary one-way ANOVA. **c**, -Thr-induced cell death can be rescued by re-addition of Thr at 8 h post starvation. Cell death (CellTox^+^ cells/confluence) was measured over time. **d**, Time course of cell death triggered by Thr starvation in the presence of z-VAD-fmk, Fer-1, NAC or Nec1s. Note that only the re-addition of Thr can rescue cell death. Cell death (CellTox^+^ cells/confluence) was measured over time by Incucyte imaging. **e**, Arg or Leu-starved cells are partly protected from the effects of Cys deprivation. Cells were deprived of Cys (black dots) or concurrent starvation with -Arg or -Leu and cell death (CellTox^+^ cells/confluence) measured over time. The data shown is representative of >10 experiments. **f**, Summary data (24 h) of the effect of Arg- or Leu-starved cells on Cys starvation-induced death from two independent experiments. **g**, ES cells were starved of Thr with concomitant withdrawal of Arg or Leu and cell death (CellTox^+^ cells/confluence) measured over time. **h**, ES cells were starved of Thr with concomitant withdrawal of each AA shown and cell death (CellTox^+^ cells/confluence) measured over time. Only concomitant loss of Met rescues cell death (boxed). **i,** ES cells were starved of Thr with concomitant withdrawal of each AA shown and cell death (CellTox^+^ cells/confluence) assessed at 24 h (n = 3). **** *p* < 0.0001 Dunnett’s multiple comparison test after ordinary one-way ANOVΑ relative to all sample groups except -Thr -Tyr.

Thr utilization is a specific and essential component of murine ES cell metabolism and is controlled by the rate-limiting enzyme threonine dehydrogenase (THD), which is not expressed in primate ES cells (and is a silenced gene in humans)^43^. In contrast to the reversible quiescence induced by Arg or Leu deprivation, Thr starvation led to complete cell death by 24 hours^43^ (**Fig. 3c**). Thr starvation-induced death took ∼8 h to initiate as re-addition of Thr at the 8 h timepoint rescued cell survival. However, this form of AA starvation death was not rescued by inhibitors of apoptosis, necroptosis or ferroptosis, suggesting the involvement of an alternative, non-conventional death pathway we provisionally term “threoptosis” (**Fig. 3d**). Thus, starvation of different AAs initiated distinct changes in cellular responses: Arg and Leu deprivation enforce cellular quiescence, while -Cys primes ES cells for ferroptosis and Thr starvation causes a non-conventional model of cell death.

### Cross-talk between AA starvation pathways

Previous results have shown that cell cycle arrest protect cells from ferroptosis, in part by preserving GSH levels^44^. We therefore hypothesized that deprivation of Arg or Leu, a process that enforces cell cycle exit, would suppress the lethal effects of Cys or Thr withdrawal. To test this idea, we simultaneously withdrew Arg or Leu in combination with Cys (i.e., -Cys, -Arg or -Cys, -Leu). Both double starvations partly blocked Cys starvation-induced ferroptosis including lipid peroxidation (**Fig. 3e, S4a-c**). However, Cys-starved ES cells eventually died, highlighting the essential nature of exogenous Cys even in quiescent cells. By contrast, withdrawal of Thr coincident with Arg or Leu could not rescue the cells. These findings further emphasize that ‘threoptosis” is a distinct form of cell death. To investigate the effects of Thr-starvation death further, we next performed a “screen” to determine if withdrawal of essential AAs beyond Arg and Leu would influence threoptosis. We created media lacking Thr combined with 13 other AAs individually dropped out (**Fig. 3h, i**). Using this pairwise approach, only withdrawal of Met rescued cell death. Importantly, the number of -Thr, -Met starved ES cells did not increase (**Fig. 3i**) but instead remained viable through 48 h of double starvation (**Fig. 3h**).

Arg or Leu starvation produced a highly transcriptional and proteomic response, including similar activation of the GCN2-ISR (**Fig. S1e, S2, S3**). We therefore wondered if Thr and Cys would also trigger similar “generic” AA starvation responses, even though the final fate of deprivation of either AA was a distinct for of cell death. We therefore investigated the transcriptional responses to Thr or Cys starvation. Surprisingly, Thr starvation massively altered the ES transcriptome with thousands of transcripts altered relative to Arg or Leu starvation combined with a heightened ISR response (**Fig. S5a**). However, genetic loss of GCN2 or inhibition of GCN2 with GCN2-IN-6 did not influence the outcome of pairwise AA dropouts and -Met rescue of threoptosis (**Fig. S6a-6**). By contrast to Thr, the absence of Cys triggered far fewer transcriptional changes compered to either -Thr or -Arg/-Leu (**Fig. S5b**). Surprisingly, the quality of the transcriptional responses was also distinct between -Cys, -Thr or -Arg/-Leu as estimated by a ranking factor metric we used to compare different RNAseq datasets (Methods) (**Fig. S5c-e**). Collectively, these results emphasize that ES cells deploy different strategies to negotiate the absence of different AAs.

### Cell cycle exit and global translational suppression confer stress protection

The effects of Met starvation on threoptosis (**Fig. 3h, i**) suggest two mutually non-exclusive possibilities to account for the death suppression: first, the absence of Met could alter the Thr catabolic pathway. Indeed, in ES cells both AAs are linked via the 1-carbon and folate cycles^43^. Second, the absence of Met could broadly affect translation at both the initiation and elongation steps. We noted that inhibition of mTOR via torin-1 but not rapamycin, which has a different mode of action^45–48^ on mTOR complexes, triggers cell cycle exit, quiescence and translation inhibition similarly to the absence of Leu or Arg (**Fig. 4a, b, Fig. 1b**) and therefore tested the latter possibility, namely that -Met mimics translation inhibition and blocks threoptosis. We compared -Met with the addition of torin-1 to Thr-starved ES cells. Remarkably, torin-1 completely rescued threoptosis (**Fig. 4c**). If translation was the critical step in this process, then a broad translation inhibitor such as cycloheximide (CHX) should produce the same effect. Indeed, CHX also blocked threoptosis (**Fig. 4d, e**). These results suggest that Thr deprivation triggers the synthesis of a protein(s) that executes non-apoptotic, -necropotic or -ferroptotic death in a Met- and mTOR-dependent manner.

**Figure 4.**
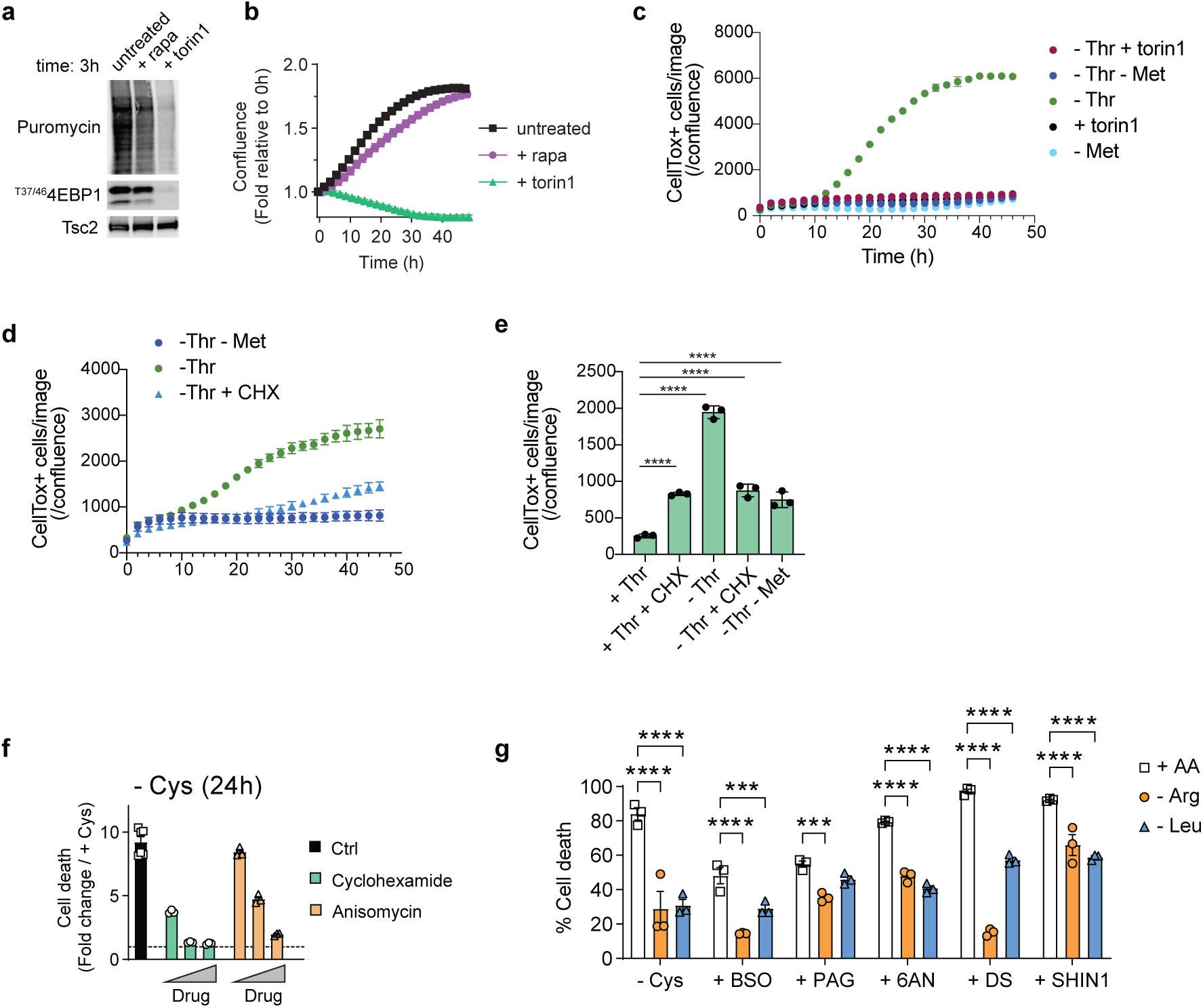
| Translational perturbation mediates protection to cell stress. **a**, Immunoblotting of translation and mTOR activity 3 h after treatment with mTOR inhibitors. Nascent proteins were labelled by puromycin. **b**, Cell proliferation in the presence of 500 nM rapamycin and torin1. Cell confluence was monitored by live imaging, normalized to 0h and shown as mean ± SEM of 3 replicates. **c**, ES cells were starved of Thr in the presence of torin-1 or concomitant withdrawal of Met and cell death (CellTox^+^ cells/confluence) measured over time. **d,** ES cells were starved of Thr in the presence of cycloheximide (CHX), or concomitant withdrawal of Met and cell death (CellTox^+^ cells/confluence) measured over time. **e,** Summary data (24 h) of the effect of Met withdrawal or CHX addition on Thr starvation-induced death (n = 3). **** *p* < 0.0001 Dunnett’s multiple comparison test after ordinary one-way ANOVA relative to the +Thr control group.**f**, Cys-starved cells are partly protected from ferroptosis by translational inhibitors CHX and anisomycin. Shown is normalized data to untreated controls (1, dotted line). The data shown is representative of >10 experiments. **g**, Cells were subjected to the following treatment for 24 h during Arg or Leu limitation: Cys deprivation, 1 mM BSO, 75 µM 6AN, 2 mM PAG, 20 µM DS or 75 µM SHIN1. Living cells were measured by flow cytometry as Annexin V-7AAD- and the percentage of cell death was calculated as the remaining cells and shown as mean ± SEM of 3 replicates. Significant differences were obtained by two-way (**d**) ANOVA followed by Dunnett’s multiple comparisons: * *p* < 0.05, ** *p* < 0.01, *** *p* < 0.001, **** *p* < 0.0001

We next investigated the effects of translation inhibition on Cys starvation. We therefore hypothesized that processes that enforce cell cycle exit via translational inhibition would also block ferroptosis in ES cells. Consistent with this idea, CHX or anisomycin also protected ES cells from Cys starvation-induced cell death (**Fig. 4f**). Collectively, therefore, cell cycle exit and translational suppression was sufficient to protect ES cells from the lethal effects of Cys withdrawal, providing a common broad protective pathway for blocking death by Cys or Thr starvation.

An emerging concept in cancer therapy is that “persister” cells that survive chemotherapy are forced, temporarily, from the cell cycle and remain in a quiescent-like state as a survival strategy ^34,49–51^. In tumor microenvironments, amino acids such as arginine and tryptophan can be limiting^52^ and as we found here for Arg and Leu, a quiescent-like state is enforced when these AAs are limiting. We therefore wondered if Arg or Leu depletion could confer broad stress protection to ES cells because starvation of these AAs enforces reversible cellular quiescence, unlike Cys or Thr starvation, which triggers cell death. We starved ES cells or Arg or Leu in combination with inhibitors of vital metabolic checkpoints including GSH synthesis, NADPH production and 1-carbon metabolism (Gclc, Cth, G6pdx, Mthfd2 or Shmt2 using L-buthionine-sulfoximine (BSO), DL-proparglyglycine (PAG), 6-aminonicotinamide (6AN), DS18561882 (DS), or SHIN1, respectively). We used -Cys as an internal control for ES cell death that is partly blocked by Arg or Leu withdrawal (**Fig. 3e, f**). As expected, all of these inhibitors were highly toxic to proliferating ES cells. However, ES cells deprived of Arg or Leu were differentially protected from the cytotoxic effects of each metabolic inhibitor (**Fig. 4h**). These results indicate that Arg or Leu deprivation, via cellular quiescence, provides partial protection against the effects of disruption of key metabolic nodes.

## Discussion

The majority of research on mammalian amino acid sensing has focused on how cells integrate amino acid availability through mTORC1, a major anabolic signaling. Most studies have used tractable, transformed cell models, such as HeLa and 293T, which rapidly respond to amino acid withdrawal by downregulating mTORC1 signalling, thereby halting anabolic growth. Further studies in various transformed cell types revealed that the routes for acquiring amino acids to support malignant growth are elastic^35,53,54^. These cells can scavenge amino acids from extracellular sources like serum and matrix^5–8,12^, by altering the expression of SLC transporters and re-wiring intrinsic pathways, including autophagy, lysosomal functions and catabolism of abundant protein complexes such as ribosomes. Moreover, the ease of manipulating amino acid signalling in these models has facilitated the identification of key amino acid-dependent upstream regulators of mTORC1, such as the GATOR complexes and their leucine and arginine regulators, Sesn2 and Castor1, respectively^13,17,18,55^.

In contrast, our study set out to map the signalling events and adaptive pathways elicited by amino acid starvation in a non-transformed cell system, aiming to understand AA signalling and adaptation in the absence of oncogene-driven pathways. We chose murine ES cells for their rapid cell division rate and experimentally reproducibility. Upon deprivation of of Leu or Arg, ES cells exit the cell cycle, reduce translation and become resistant to diverse stresses, including perturbations of glutathione and 1-carbon metabolism, which are essential for rapidly dividing cells. In contrast, when deprived of Cys or Thr, ES cells undergo cell death via distinct mechanisms: Cys deprivation induces ferroptosis while death by Thr deprivation trigger a form of cell death that is neither apoptosis, necroptosis nor ferroptosis. Notably, Thr deprivation-induced death can be rescued by co-removal of Met, or by treatment with torin-1 or other broad translation inhibitors, highlighting three distinct cellular outcomes driven by the deprivation of four amino acids. One possibility to account for the protective effects of Met withdrawal or translation inhibitors on threoptosis concerns the possibility that a newly synthesized protein(s) is necessary for the elicitation of the death pathway. Finding such a protein(s) will require a genetic-based screen and may give general insights into unexplored death pathways that intersect with core cellular metabolism.

We have limited information about how normal cells negotiate temporary restrictions in amino acids. Herein, we used the term “normal” cells as non-transformed. Further, the many types of “normal” cells in the body likely have vastly different requirements for amino acid supplies. For example, neurons are mainly non-dividing cells but have large cell masses, substantial metabolic demand and use vast amounts of some amino acids as neurotransmitter precursors (e.g., glycine, glutamine, tryptophan and tyrosine). Similarly, intestinal epithelial cells, which continuously regenerate in crypts, compete with the microbiota for nutrients, including amino acids. T cells, upon activation by antigen, rapidly alter their amino acid transporter profiles while decoupling amino acid supply from mTORC1, which responds to T cell receptor and co-stimulatory signaling^56–60^. These examples underscore the varied amino acid needs that different cell types encounter in vivo.

Our findings also indicate that forcing ES cells to exit the cell cycle is sufficient to confer protection against different metabolic stress perturbations including redox stress. Our results are consistent with recent studies in transformed cell systems where broad redox stress protection is afforded by cell cycle exit, which re-routes metabolic pathways away from translation and anabolic metabolism and toward resource conservation^44,61^. It is tempting to speculate that AA limitation may be inherently linked to cellular stress protection. In tumor microenvironments, selective AAs are limiting^52^ and arginine and tryptophan-degrading enzymes are expressed and associated with tumor survival^62,63^, which is generally thought to suppress T cell proliferation and function. In complex environments like tumors, AA starvation could affect cellular physiology in differential way: rapidly proliferating cells, including malignant cells, might be forced from the cell cycle by neighboring cells expressing AA-degrading enzymes (combined with the effects of local nutrient consumption and reduced perfusion in solid tumors). This adaptation may promote survival of some transformed cells through re-wiring of their AA response pathways. Although speculative, AA stress may protect against other forms of stress in complex in vivo milieu.

Our study has several limitations. First, ideally, a comprehensive, multi-parametric comparison of the responses to starvation by all 20 amino acids in both non-transformed and transformed cells over time would reveal distinct adaptation classes and highlight specific pathways involved in amino acid sensing and response. Such an approach should reveal “classes” of adaptation and point the way to specific pathways necessary for the detection of an AA and the corresponding adaptation to the loss of the AA. However, while the experimental scale of such an approach is feasible, the analysis of outcomes would be highly challenging. Our study showed thousands of changes in the proteomes, phospho-proteomes and transcriptomes of Arg- and Leu-starved ES cells, yet the full physiological implications of these changes remain only partially understood. Second, we did not use methods to label nascent proteins during amino acid stress^64,65^. Although translation slows following cell cycle exit, it does not cease entirely, as amino acids can still be obtained from cellular sinks even during prolonged starvation. Consequently, the identity of the proteins synthesized during complete absence of an exogenous AA remains unclear, beyond several key targets of the ISR such as ATF4 and CHOP. Future studies using unnatural AAs as labelling tools will be essential to identify and quantity nascent protein biosynthesis during amino acid limitation, potentially uncovering novel targets for addressing cancer cell persistence.

## Methods

### Cell culture

E14 mouse ES cells were a gift of Prof. D. Nedialkova (MPIB) and were originally purchased from the American Type Culture Collection (ATCC, CRL-1821). ES cells were grown on 0.1% gelatin-coated dishes in DMEM high glucose GlutaMAX containing 15% fetal bovine serum (FBS, Life Technologies), penicillin-streptomycin, β-mercaptoethanol and leukemia inhibitory factor (murine recombinant LIF, 10 ng/mL). All cell culture reagents were from ThermoFisher Scientific unless otherwise specified. All cell lines were grown in humidified tissue culture incubators at 37°C with 5% CO_2_. Cells were routinely tested to be free of Mycoplasma contamination by PCR testing (LookOut Mycoplasma PCR Detection Kit, MP0035, Sigma Aldrich).

### Single AA starvation experiments

0.5 x 10^6^ ES cells were seeded in 12-well plates for 12 h. Before starvation, cells were washed 3 times with PBS to remove any AA left. In Arg and Leu starvation experiments, we used SILAC DMEM medium (D9443, Sigma Aldrich) supplemented with penicillin-streptomycin, GlutaMAX, glucose, 5% dialyzed FBS (dialyzed in house), β-mercaptoethanol, LIF, L-Lys and with or without Arg or Leu. L-Gln starvation was performed in DMEM (D9800-13, US Biological) containing penicillin-streptomycin, 3.7 g/L sodium bicarbonate, 4.5 g/L glucose, 5% dialyzed FBS, β-mercaptoethanol, LIF and supplemented with L-Cys, Gly, L-His, L-Iso, Leu, L-Lys, L-Met, L-Phe, L-Ser, L-Trp, L-Tyr, L-Val and either Arg, Leu and/or L-Gln as indicated in the figure. AAs were from Carl Roth and made as 200X (Arg, L-His, L-Iso, Leu, L-Lys, L-Phe, L-Trp, L-Val) or 1000X (L-Cys, Gly, L-Met, L-Ser) stocks in PBS, directly resuspended in medium (L-Tyr) or obtained as a ready-made solution (L-Gln, 100X, Life Technologies).

### Drugs and other reagents

The following compounds were used as indicated in the figure legend: 6-aminonicotinamide (6AN, A68203, Sigma Aldrich), anisomycin (A9789, Sigma Aldrich), L-buthionine-sulfoximine (BSO, B2515, Sigma Aldrich, 200 mM stock resuspended in water), cyclohexamide (C7686, Sigma Aldrich), DS18561882 (DS, HY-130251, MedChemExpress), GCN2-IN-6 (resuspended in DMSO to 10 mM and used as previously described^66^), MG132 (474790, Millipore), mitomycin C (M4287, Sigma Aldrich, resuspended in water), DL-proparglyglycine (PAG, P78888, Sigma Aldrich, 400 mM stock resuspended in water), 1 μM RSL3 (S8155, SelleckChem), SHIN1 (HY-112066, MedChemExpress). Dinaciclib (HY-10492, MedChemExpress) and palbociclib (S1116, SelleckChem, resuspended in water) were used to inhibit CDK 1/2/5/9 and CDK4/6, respectively. 2 μM Ferrostatin-1 (Fe1, SML0583, Sigma Aldrich), 10 µM z-VAD-fmk (MedChemExpress in DMSO 10 mM) and 10 µM Nec1S (Cell Signaling Technologies in DMSO 10 mM) was used to inhibit ferroptosis, apoptosis and necroptosis, respectively. mTOR was directly inhibited using rapamycin (533210, Millipore) or torin1 (14379, Cell Signaling). All drugs and compounds were resuspended in DMSO unless otherwise specified.

### CRISPR/Cas9

To generate a *Eif2ak4^-/--^* genetic deficiency in ES cells, sgRNA sequences were designed using standard design tools (IDT) and cloned into the pX458 CRISPR vector (pSpCas9(BB)-2A-GFP, Addgene, #48138). Single-cell cloning was performed directly following sorting for GFP^+^ cells in 96-well plates, which were then selected based on the target protein expression and/or sequencing. The following sgRNA and crRNA sequences were used:

*Eif2ak4* sgRNA_1 (exon 9): GCAAAGTAGCGGACGATATT

*Eif2ak4* sgRNA_2 (exon 9): AGTAGCGGACGATATTTGGA

### Immunoblotting

Cell lysates were prepared on ice in RIPA buffer (ab156034, Abcam) containing protease and phosphatase inhibitors (78444, Thermo Fisher). As applicable, protein concentrations were determined using the Pierce BCA Protein Assay Kit (23225, Thermo Fisher) and a Nanodrop2000 spectrophotometer (Thermo Fisher). Proteins were separated on 4–15% Criterion TGX Stain-Free Protein Gel (5678084/5, BioRad) and transferred to 0.2 µM nitrocellulose (Amersham). Membranes were blocked in 1-3% BSA (A2153, Sigma) or 3% nonfat milk (T145.2, Carl Roth) in TBS - 0.01% Tween 20 and probed overnight at 4°C with primary antibodies listed below. Membranes were washed and probed with secondary antibodies and developed using SuperSignal West Pico PLUS Chemiluminescent reagent (34580, Thermo Fisher). Homogenous loading was controlled by blotting for Vinculin, Grb2 or Tsc2.

### Antibodies

The following antibodies were used in immunoblotting experiments: phospho-4EBP1 (T37/46, 1:1,000, 2855, Cell Signaling), Atf4 (1:1,000, sc-390063, Santa Cruz), phospho-Gcn2 (T899, ab75836, 1:1000, Abcam), Grb2 (1:1,000, 610112, BD Biosciences), puromycin (1:10,000, MABE343, Millipore), phospho-S6K (T389, 1:1,000, 9234, Cell Signaling), Tsc2 (1:1,000, 4308, Cell Signaling), Vinculin (1:2000, 66305-1-Ig, Proteintech).

### Estimation of mRNA translation

To estimate translation rates, we used surface sensing of translation (SUnSET) method^67^ where puromycin incorporation in neosynthesized proteins approximates the rate of mRNA translation at a given time point. 10 µg/ml of puromycin (P8833, Sigma Aldrich, 10 mg/ml stock resuspended in PBS) was added to the culture 10 min before sample collection. Samples were then subjected to western blotting. When indicated, the following drugs were used to inhibit translation: cycloheximide (C7698, Sigma Aldrich).

### Cell proliferation and cell death analyses

E14 ES cells were seeded in 48-well plates and treated as specified in the figure legend. Non-pharmacological ferroptosis induction was performed using L-Cys-free DMEM (D9800-13, US Biological) containing penicillin-streptomycin, 3.7 g/L sodium bicarbonate, 4.5 g/L glucose, 10% dialyzed FBS, β-mercaptoethanol, LIF and the relevant AAs as described above. To evaluate proliferation, cells were fixed and permeabilized using BD Cytofix/Cytoperm and stained with anti-Ki67 FITC antibody (652410, Biolegend). All reagents were from BD Biosciences unless specified. All measurements were recorded on the LSR Fortessa with the FACS Diva software (BD Biosciences) and further analyzed using FlowJo 10 software. *Live microscopy.* Data acquisition and analysis were performed using the IncuCyte^®^ S3 Live Cell Analysis System (Sartorius, USA). Nine images per replicate were taken every 2h for 48h using the 10X objective. Cell proliferation was evaluated by analyzing the cell confluence in the phase channel. For cell death measurements, CellTox Green (G8731, Promega) was used according to the manufacturer’s instructions to stain cells with impaired membrane integrity. CellTox^+^ green object numbers were then counted in the fluorescence channel and normalized to the cell confluence for each well. The results are shown as a percentage of cell death after further normalization against the unstarved group in each dataset using 0% cell death as the 0h time point and 100% as the maximum cell death seen during the time course.

### Measurement of ROS production, lipid peroxidation and mitochondrial potential

ROS production and lipid peroxidation were assessed according to the manufacturer’s instructions. Briefly, 10 μM of CM-H_2_DCFDA (C6827, Thermo Fisher) or 2 μM of C11-BODIPY 581/591 (D3861, Thermo Fisher) or 20 nM of Tetramethylrhodamine Methyl Ester (TMRM, M20036, Thermo Fisher) was added to the cells 30 min before the end of the experiment. 20 μM of Carbonyl cyanide 3-chlorophenylhydrazone (CCCP) was added to the cells for 5 min to induce mitochondrial membrane depolarization as a control. The MFI in the samples were measured by flow cytometry (LSR Fortessa, BD) and analyzed using FlowJo 10 software.

### RNAseq

Cells were starved of Arg, Leu versus control unstarved ES cells or Cys or Thr versus unstarved cells in an independent experiment set, all in triplicates. RNA was extracted using Direct-zol RNA Miniprep Plus (R2071, Zymo Research) according to the manufacturer’s protocol. mRNA sequencing libraries were prepared with 1 µg of total RNA of each sample using the NEBNext Ultra II Directional RNA Library Prep Kit for Illumina (E7765, NEB) with NEBNext Poly(A) mRNA Magnetic Isolation Module (E7490, NEB), according to standard manufacturer’s protocol. Total RNA and the final library quality controls were performed using Qubit Flex Fluorometer (Q33327, Thermo Fisher Scientific) and 2100 Bioanalyzer Instrument (G2939BA, Agilent) before and after library preparation. Paired-end sequencing was performed on Illumina NextSeq 500 (2 × 43 bp reads). The samples were multiplexed and sequenced on one High Output Kit v2.5 to reduce batch effects. BCL raw data was converted to FASTQ data and demultiplexed by bcl2fastq Conversion Software (Illumina). After checking the quality of the samples (FastQC, v.0.11.7), the files were mapped to the mouse genome (Genome build GRCm38) downloaded from Ensembl using the star aligner^68^ version 2.7.4a. The mapped files were then quantified on a gene level based on the Ensembl annotations, using the HTseq tool^69^ in Python. Using the DESeq2 package^70^ (R 4.0.3, DESeq version 1.30.1), the count data was normalized by the size factor to estimate the effective library size. A filtering step of removing genes with less than 1 reads was used. This followed by the calculation of gene dispersion across all samples. The analysis of two different conditions against each other resulted in a list of differentially expressed genes for each comparison. Genes with an adjusted p-value of ≤ 0.05 were then considered to be differentially expressed for downstream analysis.

### Comparison of RNAseq datasets

To enable a meaningful comparison of datasets, we use a customized Ranking Factor (RF), defined as:

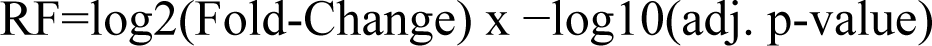

This approach combines both the magnitude of gene expression change (log2 fold-change) and the statistical significance (adjusted p-value) into a single metric, providing a more robust measure of gene importance. By integrating these two factors, the RF prioritizes genes that exhibit both substantial expression changes and strong statistical support, ensuring that biologically relevant genes are highlighted based on both their expression level and statistical confidence.

### Phosphoproteomics and total proteomics

*Sample preparation for MS analysis.* All MS experiments were performed in biological quadruplicates. E14 ES cells were grown on 10 cm dishes and starved of Arg or Leu for 1, 3, or 10 h. A time point 0h (unstarved) was used as a control. Cells were then washed three times with ice-cold TBS, lysed in 4% sodium deoxycholate (SDC, 30970, Sigma Aldrich) and 100 mM Tris-HCl pH 8.5 (T6066, Sigma Aldrich), and boiled immediately. After sonication, protein amounts were quantified (BCA) and 1.1 mg for each sample was used for digestion. Samples were reduced with 10 mM tris(2-carboxy(ethyl)phosphine (TCEP), alkylated with 40 mM 2-chloroacetamide (CAA) and digested with trypsin and lysC (1:100, enzyme/protein, w/w) overnight. For proteome measurements, 20 µg of the total peptide was desalted using SDB-RPS stage tips. 500 ng of desalted peptides were resolubilized in 5 µl 2% ACN and 0.3% TFA and injected into the mass spectrometer. To enrich phosphorylated peptides, we applied the EasyPhos protocol previously described^71^. In short, a solution of 50% isopropanol, 6% trifluoroacetic acid (TFA) and 1 mM monopotassium phosphate (KH2PO4) was added to the digested lysate. Lysates were shaken, then spun down for 3 min at 2000×g, and supernatants were incubated with TiO_2_ beads for 5 min at 40 °C (1:10, protein/beads, w/w). Beads were washed 5 times with isopropanol and 5% TFA, and phosphopeptides were eluted with 40% acetonitrile (ACN) and 15% of ammonium hydroxide (25% NH_4_OH) on C8 stage tips. After 20 min of SpeedVac at 45 °C, phosphopeptides were desalted on SDB-RPS stage tips and resolubilized in 5 µl 2% ACN and 0.3% TFA and injected in the mass spectrometer. *LC-MS.* Samples were loaded onto 50-cm columns packed in-house with C18 1.9 μM ReproSil particles (Dr. Maisch GmbH), with an EASY-nLC 1000 system (Thermo Fisher Scientific) coupled to the MS (Q Exactive HFX, Thermo Fisher Scientific). A homemade column oven maintained the column temperature at 60°C. Peptides were introduced onto the column with buffer A (0.1% formic acid), and phosphopeptides were eluted with a 70 min gradient starting at 3% buffer B (80% ACN, 0.1% formic acid) and followed by a stepwise increase to 19% in 40 min, 41% in 20 min, 90% in 5 min and 95% in 5 min, at a flow rate of 300 nL/min, while peptides for proteome analysis were eluted with a 120 min gradient starting at 5% buffer B (80% ACN, 0.1% formic acid) followed by a stepwise increase to 30% in 95 min, 60% in 5 min, 95% in 2 x 5 min and 5% in 2 x 5 min at a flow rate of 300 nL/min. A data-independent acquisition MS method was used for proteome and phosphoproteome analysis in which one full scan (300 to 1650 *m/z*, R = 60,000 at 200 *m/z*) at a target of 3 × 10^6^ ions was first performed, followed by 32 windows with a resolution of 30,000 where precursor ions were fragmented with higher-energy collisional dissociation (stepped collision energy 25%, 27.5%, 30%) and analyzed with an AGC target of 3 × 10^6^ ions and a maximum injection time at 54 ms in profile mode using positive polarity. *MS data analysis.* MS raw files were processed by the Spectronaut software version 13 (Biognosys) using the mouse Uniprot FASTA database (22,220 entries, 39,693 entries, 2015). Proteome and phosphoproteome files were analyzed via direct DIA. For proteome analysis, standard settings were used. The false discovery rate (FDR) was set to less than 1% at the peptide and protein levels and a minimum length of 7 AAs for peptides was specified. Enzyme specificity was set as C-terminal to Arg and L-Lys as expected using trypsin and LysC as proteases and a maximum of two missed cleavages. For the phosphoproteome analysis, serine/threonine/tyrosine phosphorylation was added as variable modification to the default settings which include cysteine carbamidomethylation as fixed modification and N-terminal acetylation and methionine oxidations as variable modifications. The localization cutoff was set to 0. *Bioinformatics data analysis.* We performed downstream analyses in the Perseus software version 1.6.2.2_72_. For phosphosite analysis, Spectronaut normal report output tables were collapsed to phosphosites and the localization cutoff was set to 0.75 using the peptide collapse plug-in tool for Perseus as previously described^73^, which collapses phosphoions to phosphosites. Importantly, it does not sum up the intensities of a phosphosite on peptides, if different phosphorylations are also present. For example, the intensity of Larp1_S743_M1 and Larp1_S743_M2 differs because while Larp1_S743_M1 represents the singly phosphorylated peptide, Larp1_S743_M2 reflects two phosphorylated sites on one (or more) peptides containing S743 (Larp1_S743_S751 and Larp1_S743_T747 in this case). For each phosphosite on a multiple phosphorylated peptide, we receive a row with the same intensities as these phosphorylations are localized on the same peptide. Eif4ebp1_S64_M2 and Eif4ebp1_T69_M2 share the same intensity as they represent the two phosphosites on the same peptide. Different phosphosites on the same peptide can have slightly different fold changes due to imputation. Also, each collapse key (gene_position_multiplicity) is unique, which means that if a phosphosite is present on two peptides that carry a different phosphosite, just one row will be assigned. Phosphosites located on phosphopeptides with more than two phosphorylations are labeled with a multiplicity of 3 (M3). Summed intensities were log_2_ transformed. Quantified proteins and phosphosites were filtered for at least 75% of valid values among three or four biological replicates in at least one condition. Missing values were imputed and significantly up-or down-regulated hits were determined by multiple-sample test (one-way analysis of variance (ANOVA), *s0* = 0.1, FDR = 0.05) and Student’s t-test (two-sided), (*s0* = 0.1, FDR = 0.05). For hierarchical clustering of significant proteins and phosphosites, median abundances of biological replicates were z-scored and clustered using Euclidean as a distance measure for row clustering. Fisher’s exact tests were performed to detect the systematic enrichment of annotations and pathways by analyzing proteins whose levels or expression or phosphorylation levels are significantly regulated upon different conditions. We used KEGG names, motifs and PhosphoSitePlus (PSP) kinase/substrate annotations and either the *p*-value or the Benjamini-Hochberg FDR was set to 0.02 as a threshold. The PSP database contains 2171 known mouse kinases/substrates annotations (including 87 mTOR kinase/substrates). The identity and position of the phosphorylation sites on 4ebp1 and Ulk1 were inferred from PSP.

### Seahorse bioenergetic measurements

25,000 ES cells were plated into each well of Seahorse cell culture plates pre-coated with 0.1% gelatin. 5h later, starvation experiments were started as described above and carried out overnight (17h). Real-time oxygen consumption rate measurements were made with a Seahorse XF HS Mini Analyzer (Agilent) and the Seahorse XFp Cell Mito Stress kit (103010-100, Agilent) following the manufacturer’s instructions. Briefly, before running the assay, the plates were preincubated at 37°C for a minimum of 45 min in the absence of CO_2_ in Seahorse XF DMEM medium (103575-100; Agilent) supplemented with 2 mM glutamine, 10 mM glucose, and 1 mM pyruvate. Cells were stimulated with 1.5 μM oligomycin, 1 μM Carbonyl cyanide-4 (trifluoromethoxy) phenylhydrazone (FCCP) and 0.5 μM AntimycinA/Rotenone. After the experiment, cells were lysed in RIPA and the protein amounts quantified in each sample to allow normalization between conditions.

### Targeted metabolomics

*Sample preparation.* All samples were prepared in biological triplicates. ES cells were starved of Arg or Leu for 1, 3 and 10 hours as described before and medium containing all amino acids was used as a control. After the indicated time points, cells were washes with PBS and lysed using a pre-cooled MeOH:ACN:H_2_O (2:2:1, v/v) solvent mixture (MeOH and ACN were ACS grade and from Sigma Aldrich and AppliChem, respectively). Cell lysates were then vortexed for 30 s, and incubated in liquid nitrogen for 1 min. The samples were then allowed to thaw at room temperature and sonicated for 10 min in a water bath sonicator. The cycle of cell lysis in liquid nitrogen combined with vortex and sonication was repeated two times. To precipitate proteins, the samples were incubated for 1h at −20 °C, followed by a 15 min centrifugation at 13,000 rpm at 4 °C. After centrifugation, the supernatants were snap frozen in liquid nitrogen, and stored at −80 °C. *LC-MS.* Metabolite extracts have been analyzed by either reversed phase chromatography or hydrophilic interaction chromatography (HILIC), both directly coupled to mass spectrometry (LC-MS/MS). For reversed phase LC-MS/MS, 100 µl of the extracts have been dried down in a vacuum centrifuge and resolved in 100 µl of 0.1% formic acid in water. For each sample, 1 µl has been injected onto a Kinetex (Phenomenex) C18 column (100 Å, 150 x 2.1 mm) connected with the respective guard column, and employing a 7-minute-long linear gradient from 99% A (1 % acetonitrile, 0.1 % formic acid in water) to 60% B (0.1 % formic acid in acetonitrile) at a flow rate of 80 µl/min. Detection and quantification has been done by LC-MS/MS, employing the selected reaction monitoring (SRM) mode of a TSQ Altis mass spectrometer (Thermo Fisher Scientific), using the following transitions in the positive ion mode: 76 m/z to 30 m/z (glycine), 90 m/z to 44 m/z (alanine), 106 m/z to 60 m/z (serine), 116 m/z to 70 m/z (proline), 118 m/z to 72 m/z (valine), 120 m/z to 74 m/z (threonine), 122 m/z to 76 m/z (cysteine), 132 m/z to 86 m/z (leucine and isoleucine),133 m/z to 74 m/z (asparagine), 134 m/z to 74 m/z (aspartic acid), 147 m/z to 84 m/z (lysine), 147 m/z to 130 m/z (glutamine), 148 m/z to 84 m/z (glutamic acid), 150 m/z to 133 m/z (methionine), 156 m/z to 110 m/z (histidine), 166 m/z to 133 m/z (phenylalanine), 175 m/z to 70 m/z (arginine), 182 m/z to 36 m/z (tyrosine), 205 m/z to 188 m/z (tryptophan), 241 m/z to 74 m/z (cystine). In HILIC, 1 µl of the original sample was directly injected onto a polymeric iHILIC-(P) Classic HPLC column (HILICON, 100 x 2.1 mm; 5 µm) and the respective guard column, operated at a flow rate of 100 µl/min. The HPLC (Ultimate 3000 HPLC system; Dionex, Thermo Fisher Scientific) was directly coupled via electrospray ionization to a TSQ Quantiva mass spectrometer (Thermo Fisher Scientific). A linear gradient (A: 95% acetonitrile 5%, 10 mM aqueous ammonium acetate; B: 5 mM aqueous ammonium bicarbonate) starting with 15% B and ramping up to 60% B in 9 minutes was used for separation. The following SRM transitions were used for quantitation in the negative ion mode: 87 m/z to 43 m/z (pyruvate), 140 m/z to 79 m/z (carbamoyl phosphate), 191 m/z to 111 m/z (citrate), 229 m/z to 97 m/z (pentose phosphates), 259 m/z to 97 m/z (hexose phosphates), 426 m/z to 134 m/z (ADP), 506 m/z to 159 m/z (ATP), 662 m/z to 550 m/z (NAD), 664 m/z to 408 m/z (NADH), 742 m/z to 620 m/z (NADP), 744 m/z to 426 m/z (NADPH), 808 m/z to 408 m/z (acetyl-CoA). *Data analysis.* Data interpretation was performed using TraceFinder (Thermo Fisher Scientific). Authentic standards were used for determining collision energies and for validating experimental retention times via standard addition. The metabolite peaks from raw HPLC-MS chromatograms were integrated in the software by evaluating the extracted ion chromatogram (XIC) counts. All data sets were normalized to the amount of protein in the samples (determined by the Pierce BCA protein assay kit, 23227, Thermo Fisher) to account for cell growth. Data is shown as XIC area/ug of protein, log_2_ transformed result as indicated in the figure legends.

### Statistics and reproducibility

The statistical analyses shown in bar plots were performed with GraphPad Prism 9 using one-way or two-way ANOVA followed by Dunnett’s test for multiple comparisons, unless otherwise mentioned. A *p*-value less than 0.05 was considered significant. All non-omics experiments were independently repeated more than two times with similar results. Omics-related statistical analyses are described in the previous sections above.

## Supporting information

RNAseq data -Leu -Arg

Phosphoproteomic data -Leu -Arg

Proteomic data -Leu -Arg

RNAseq data -Cys -Thr

## Data availability

All the MS data generated for this study have been deposited to the ProteomeXchange Consortium via the PRIDE repository (http://www.proteomexchange.org/) with the identifier PXD031093. RNAseq data have been deposited in the Gene Expression Omnibus repository (https://www.ncbi.nlm.nih.gov/geo/) under the accession numbers GSE195470 and GSE291532. All the remaining data are available within the Article or from the author upon reasonable request. The MPIB NGS Core facility (RRID:SCR_025746) and MPIB Bioinformatics Core facility (RRID: SCR_025742) were used for this project.

## Code availability

The manuscript does not report any custom code.

### Acknowledgments

We thank Professors Brenda Schulman and Danny Nedialkova at MPIB for reagents, Sabine Suppmann and the MPIB Protein Production Core Facility for LIF. We also acknowledge Dr. Rin Ho Kim and the MPIB NGS core facility for RNAseq sample preparation, data acquisition and processing. Dr. Thomas Köcher and the Vienna BioCenter Core Facilities (VBCF, Vienna, Austria) for metabolomics sample preparation, data acquisition and processing. The VBCF Metabolomics Facility is funded by the City of Vienna through the Vienna Business Agency. We thank Hiromune Eto for helping with RNAseq analysis and Leo Kiss, Leonie Zeitler, Johanna Brüggenthies and Patricia Ogger for scientific input. Work in the Murray laboratory is supported by the Max-Planck-Gesellschaft.

## Author contributions

M.R. and P.J.M. conceived the study and wrote the paper. M.R. performed all the experiments unless otherwise indicated and analysed the data. A. F. performed the cell death flow cytometry experiment. A.G. helped with the Seahorse experiment. A. F. and A.G. contributed to methods development and discussed the results. A.Y. performed bioinformatic analysis of RNAseq data. M. T. performed the total and phosphoproteomics sample preparation, data acquisition and helped with MS data analysis. M.M. provided MS expertise. All authors reviewed the manuscript.

## Supplemental figures

**Figure S1.**
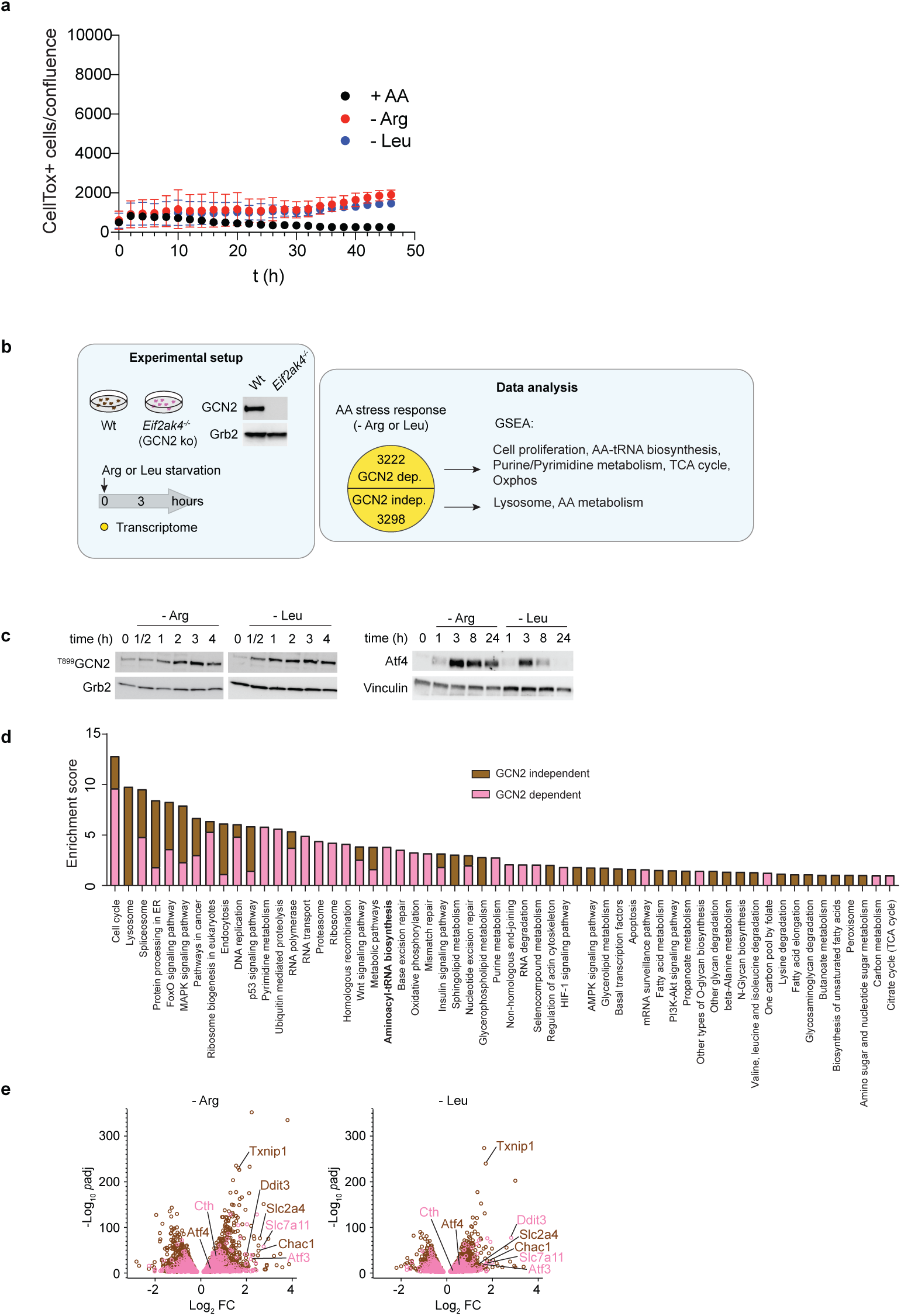
**a**, Withdrawal of Arg or Leu does not induce cell death. Shown are the ratio of CellTox+ cells to the confluence across a time window of 48 hr. **b**, Design of the RNAseq experiment to compare control and GCN2-deficient ES cells. Validation of the complete loss of GCN2 is shown in the left inset. **c,** Immunoblotting of the induction of GNC2 response and the ISR shortly after Arg or Leu deprivation. **d**, Enrichment analysis was performed on significantly regulated transcripts obtained in both transcriptomes using *Mus musculus* KEGG annotations and DAVID databases. Top enriched pathways are shown (FDR < 5%) and ranked by their fold enrichment. The hits are color-coded regarding their GCN2 dependency: in brown, the genes that are significantly regulated in the GCN2 knock-out cells (GCN2 independent), in pink, the genes that are not significantly regulated in the GCN2 knock-out cells (GCN2 dependent). **e,** Volcano plots representing statistical significance (-log_10_ *p*-value) versus magnitude of changes (log_2_ FC) of all significantly regulated genes in Wt cells at 3 hr Arg or Leu starvation. The hits are color-coded regarding their GCN2 dependency: in brown, the genes that are significantly regulated in the GCN2 knock-out cells (GCN2 independent), in pink, the genes that are not significantly regulated in the GCN2 knock-out cells (GCN2 dependent).

**Figure S2.**
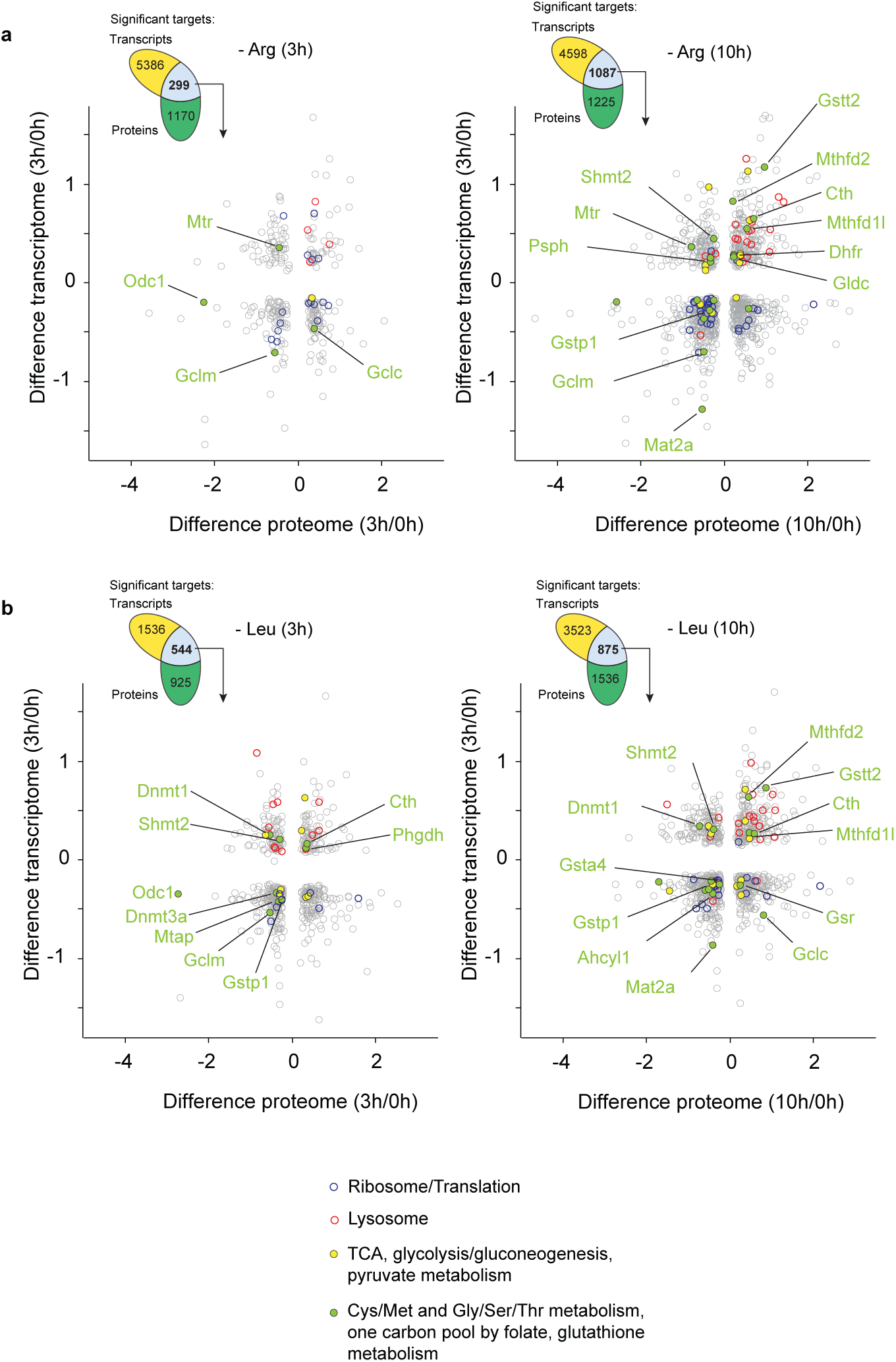
**a, b,** Correlations of proteome and transcriptome differences (log_2_ FC starvation/unstarved groups) during Arg or Leu deprivation. All common significantly regulated transcripts and proteins are shown. Selected proteins annotated with KEGG or GO terms are color-coded: in blue, 03010 and GOBP 0006412; in red, 04142; in yellow, 00010, 00020, 00620; in light green: 00260, 00270, 00480, 00670.

**Figure S3.**
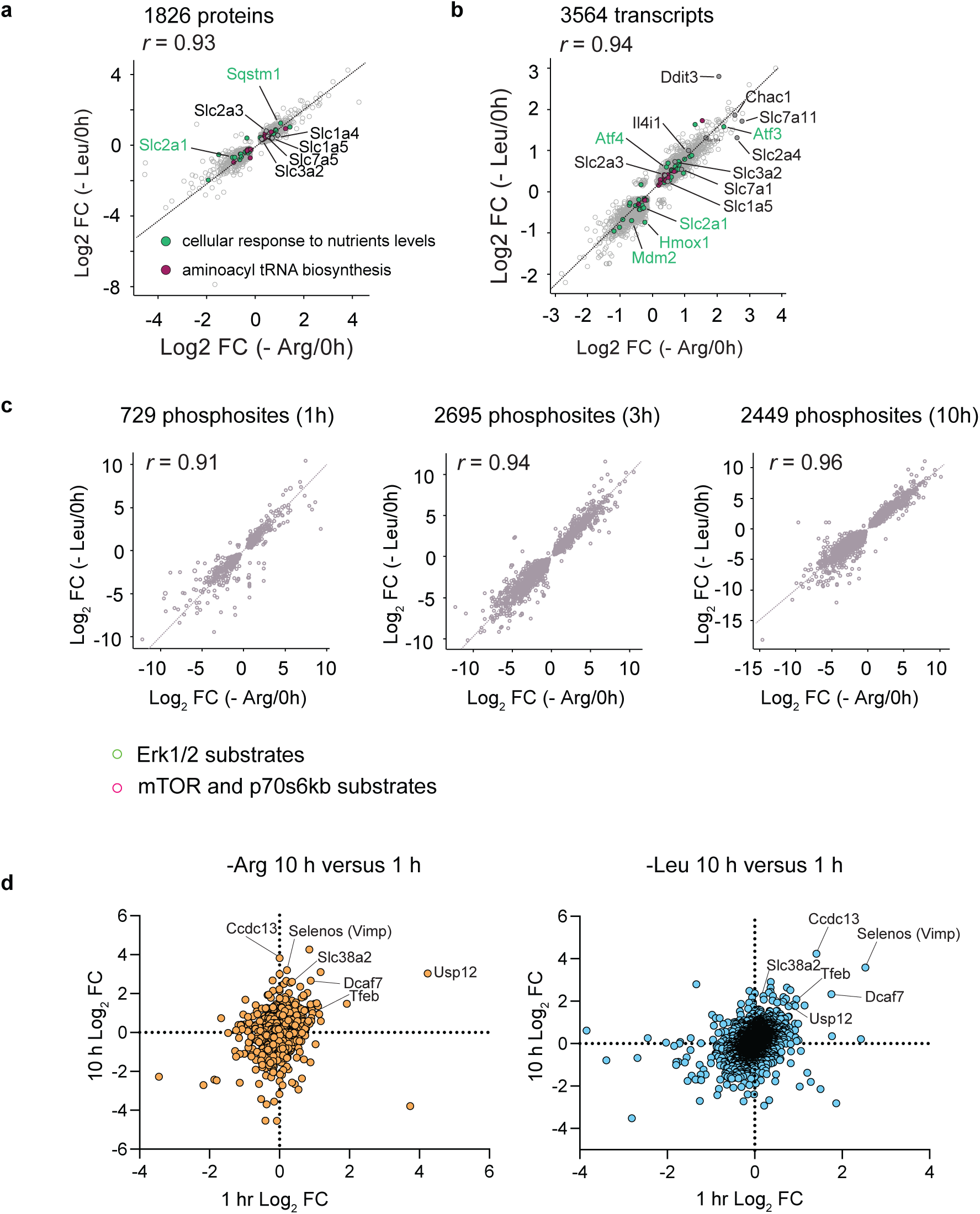
**a, b**, Spearman correlations (two-tailed *p* < 0.0001) of Arg and Leu starvation response in the proteome (r = 0.93) and transcriptome (r = 0.94). Selected targets such as nutrient transporters are colored according to GOBP and KEGG annotations: in green, 0031669; in purple, 00970. **c**, Spearman correlations of Arg and Leu starvation response in the phosphoproteome at the indicated time points (two-tailed *p* < 0.0001). All Spearman correlation analyses were performed on common significantly regulated transcripts and proteins in both Arg and Leu groups. **d,** Volcano plots showing transcript variation with Arg or Leu starvation at 3 h. Representative common (comparing -Arg versus -Leu) transcripts increased in the starvation condition are annotated.

**Figure S4.**
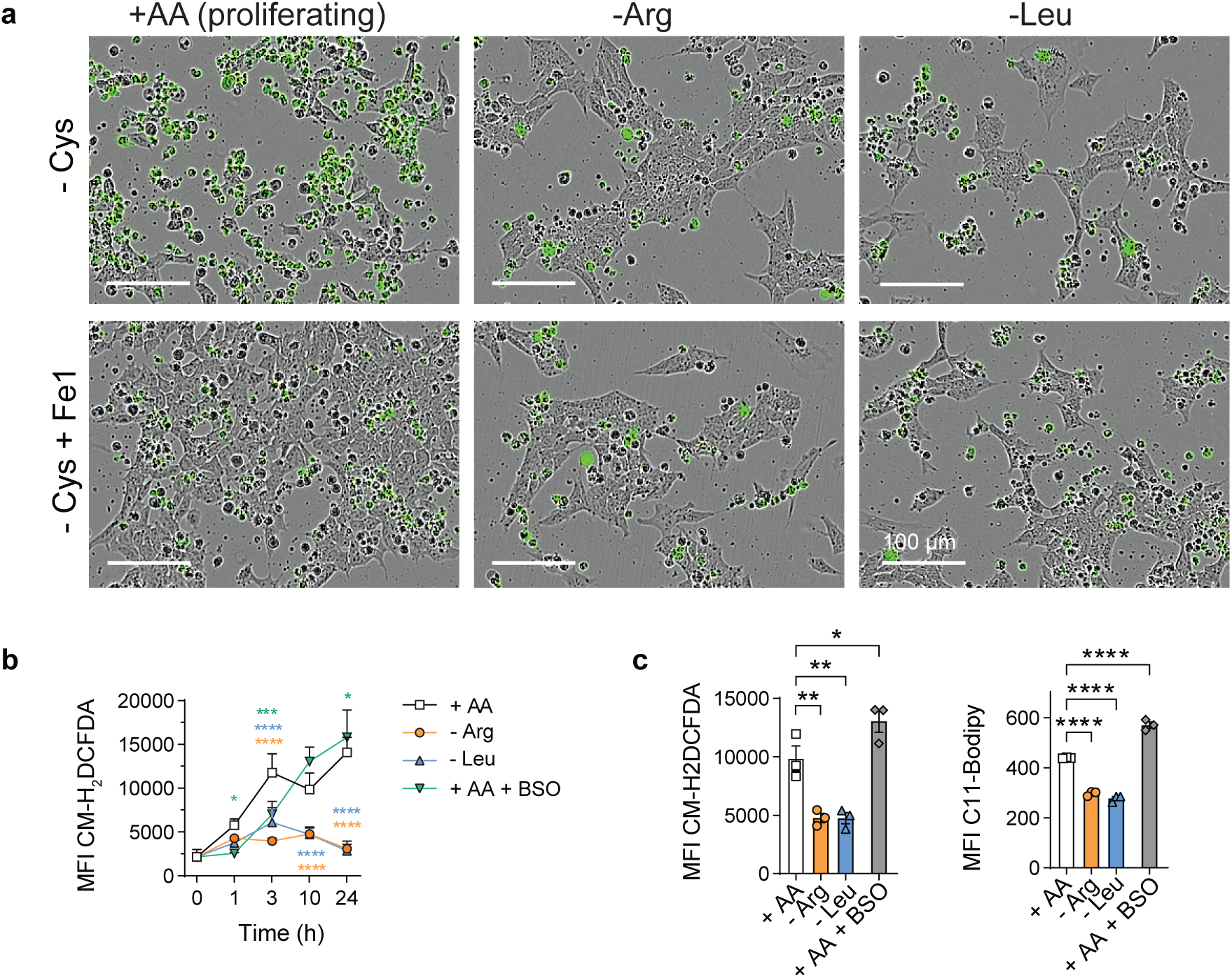
**a**, Cys deprivation in the presence or absence of Arg or Leu. Representative images are shown and dead cells are CellTox+ green objects. As a control (lower images) 2 μM Fe1 was added to inhibit ferroptosis. **b**, Time course of intracellular levels of ROS in -Arg, -Leu or proliferating control cells (+AA) was quantified using the CM-H2DCFDA fluorescent probe. As a positive control for ROS production, cells in complete media were treated with 100 μM BSO (+AA +BSO). Data is depicted as MFI ± SEM of 3 replicates and significant differences were obtained by two way-ANOVA followed by Dunnett’s multiple comparisons at each time point: * p < 0.05, *** p < 0.001, **** p < 0.0001. **c**, Single timepoint measurements of intracellular levels of ROS (left panel) or lipid peroxidation (right panel) was quantified 10 h or 24 h later using the CM-H2DCFDA or C11-Bodipy fluorescent probes, respectively. As a control, cells in normal media were treated with 100 μM BSO. Data is depicted as the MFI of the mean ± SEM of 3 replicates.

**Figure S5.**
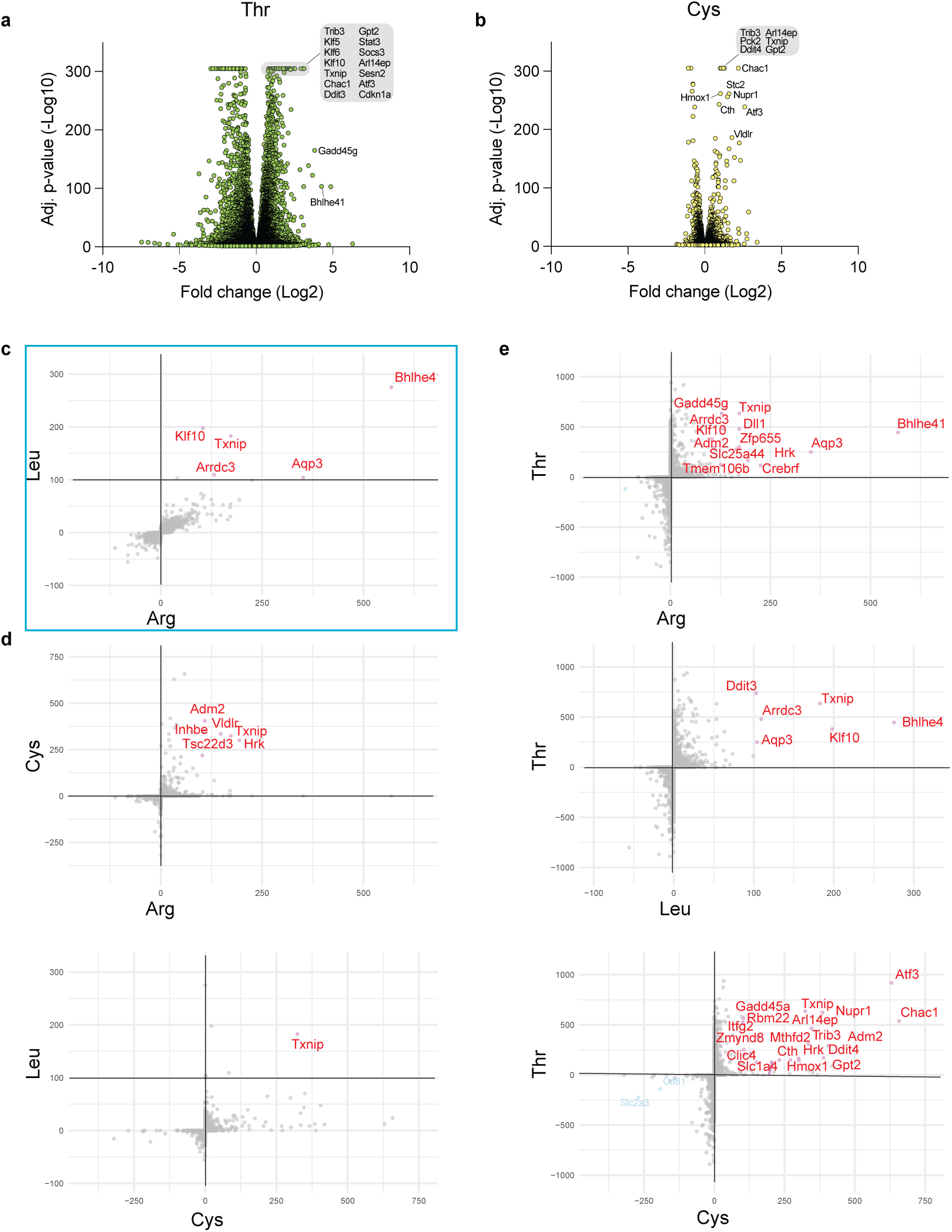
**a, b**, Volcano plots showing transcript variation with Thr or Cys starvation at 3 h. **c**, Correlation between Leu and Arg transcriptional responses at 3 h shown by ranking factor (Methods). **d, e** Correlation between Thr and Leu or Thr and Arg or Cys and Thr transcriptional responses at 3 h.

**Figure S6.**
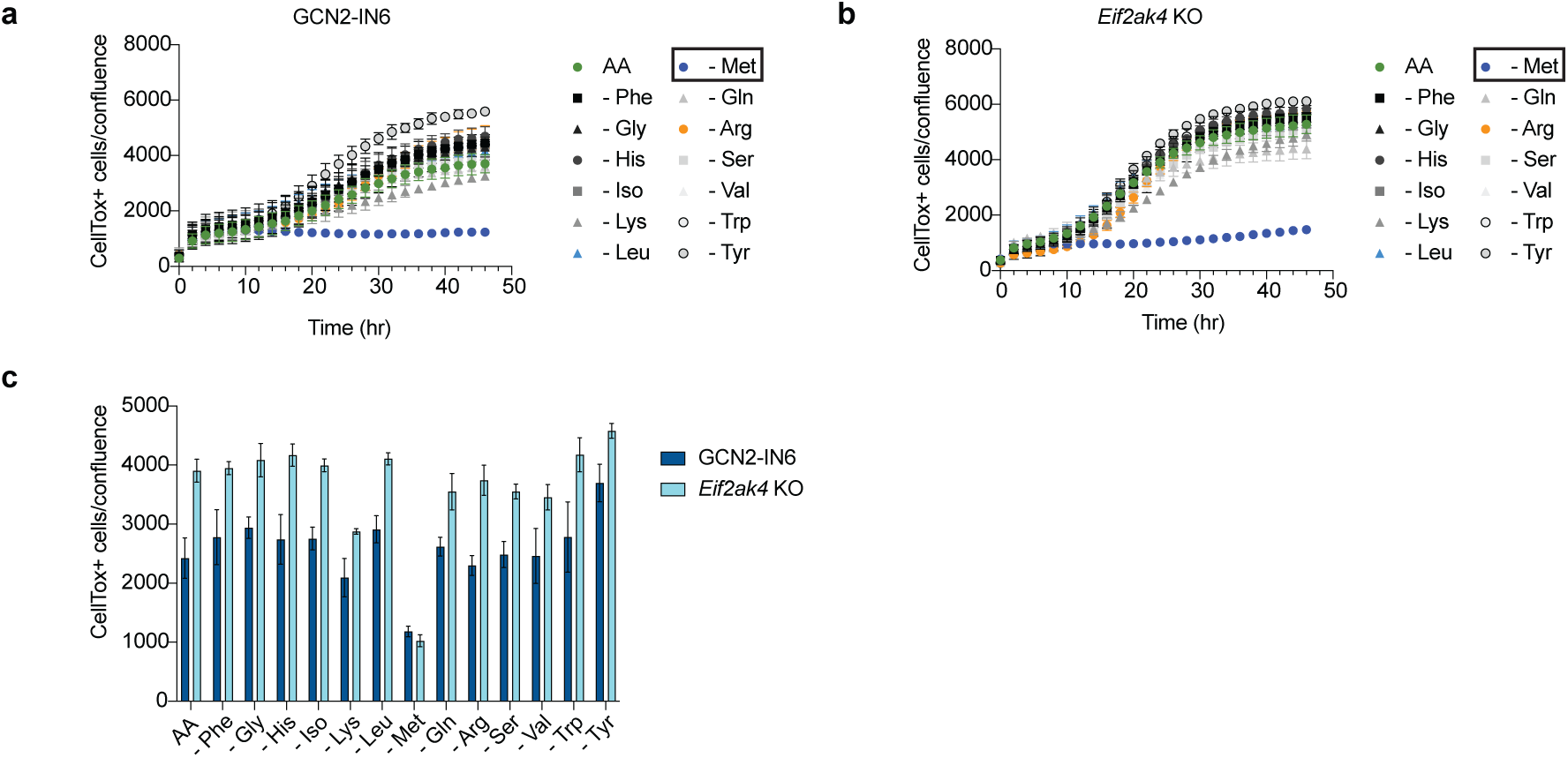
**a,b**, Rescue of -Thr-induced cell death by concomitant loss of Met is independent of GCN2 as shown by the addition of GCN2-IN-6 or genetic disruption of *Eif2ak4* (encoding GCN2). **c**, Summary data at 24 h for panels c, d.

## References

1. Gu, X., Jouandin, P., Lalgudi, P.V., Binari, R., Valenstein, M.L., Reid, M.A., Allen, A.E., Kamitaki, N., Locasale, J.W., Perrimon, N., and Sabatini, D.M. (2022). Sestrin mediates detection of and adaptation to low-leucine diets in Drosophila. Nature 608, 209–216. 10.1038/s41586-022-04960-2.

2. Morris, J.G., and Rogers, Q.R. (1978). Ammonia intoxication in the near-adult cat as a result of a dietary deficiency of arginine. Science 199, 431–432. 10.1126/science.619464.

3. Zhu, J., and Thompson, C.B. (2019). Metabolic regulation of cell growth and proliferation. Nat Rev Mol Cell Biol 20, 436–450. 10.1038/s41580-019-0123-5.

4. Armenta, D.A., Laqtom, N.N., Alchemy, G., Dong, W., Morrow, D., Poltorack, C.D., Nathanson, D.A., Abu-Remalieh, M., and Dixon, S.J. (2022). Ferroptosis inhibition by lysosome-dependent catabolism of extracellular protein. Cell Chem Biol. 10.1016/j.chembiol.2022.10.006.

5. Commisso, C., Davidson, S.M., Soydaner-Azeloglu, R.G., Parker, S.J., Kamphorst, J.J., Hackett, S., Grabocka, E., Nofal, M., Drebin, J.A., Thompson, C.B., et al. (2013). Macropinocytosis of protein is an amino acid supply route in Ras-transformed cells. Nature 497, 633–637. 10.1038/nature12138.

6. Kamphorst, J.J., Nofal, M., Commisso, C., Hackett, S.R., Lu, W., Grabocka, E., Vander Heiden, M.G., Miller, G., Drebin, J.A., Bar-Sagi, D., et al. (2015). Human pancreatic cancer tumors are nutrient poor and tumor cells actively scavenge extracellular protein. Cancer Res 75, 544–553. 10.1158/0008-5472.CAN-14-2211.

7. Nofal, M., Wang, T., Yang, L., Jankowski, C.S.R., Hsin-Jung Li, S., Han, S., Parsons, L., Frese, A.N., Gitai, Z., Anthony, T.G., et al. (2022). GCN2 adapts protein synthesis to scavenging-dependent growth. Cell Syst 13, 158–172 e159. 10.1016/j.cels.2021.09.014.

8. Nofal, M., Zhang, K., Han, S., and Rabinowitz, J.D. (2017). mTOR Inhibition Restores Amino Acid Balance in Cells Dependent on Catabolism of Extracellular Protein. Mol Cell 67, 936–946 e935. 10.1016/j.molcel.2017.08.011.

9. Palm, W., Park, Y., Wright, K., Pavlova, N.N., Tuveson, D.A., and Thompson, C.B. (2015). The Utilization of Extracellular Proteins as Nutrients Is Suppressed by mTORC1. Cell 162, 259–270. 10.1016/j.cell.2015.06.017.

10. Guzelsoy, G., Elorza, S.D., Ros, M., Schachtner, L.T., Hayashi, M., Hobson-Gutierrez, S., Rundstrom, P., Brunner, J.S., Pillai, R., Walkowicz, W.E., et al. (2025). Cooperative nutrient scavenging is an evolutionary advantage in cancer. Nature. 10.1038/s41586-025-08588-w.

11. Nazemi, M., Yanes, B., Martinez, M.L., Walker, H.J., Pham, K., Collins, M.O., Bard, F., and Rainero, E. (2024). The extracellular matrix supports breast cancer cell growth under amino acid starvation by promoting tyrosine catabolism. PLoS Biol 22, e3002406. 10.1371/journal.pbio.3002406.

12. Sousa, C.M., Biancur, D.E., Wang, X., Halbrook, C.J., Sherman, M.H., Zhang, L., Kremer, D., Hwang, R.F., Witkiewicz, A.K., Ying, H., et al. (2016). Pancreatic stellate cells support tumour metabolism through autophagic alanine secretion. Nature 536, 479–483. 10.1038/nature19084.

13. Chantranupong, L., Scaria, S.M., Saxton, R.A., Gygi, M.P., Shen, K., Wyant, G.A., Wang, T., Harper, J.W., Gygi, S.P., and Sabatini, D.M. (2016). The CASTOR Proteins Are Arginine Sensors for the mTORC1 Pathway. Cell 165, 153–164. 10.1016/j.cell.2016.02.035.

14. Costa-Mattioli, M., and Walter, P. (2020). The integrated stress response: From mechanism to disease. Science 368. 10.1126/science.aat5314.

15. Saxton, R.A., Chantranupong, L., Knockenhauer, K.E., Schwartz, T.U., and Sabatini, D.M. (2016). Mechanism of arginine sensing by CASTOR1 upstream of mTORC1. Nature 536, 229–233. 10.1038/nature19079.

16. Saxton, R.A., Knockenhauer, K.E., Wolfson, R.L., Chantranupong, L., Pacold, M.E., Wang, T., Schwartz, T.U., and Sabatini, D.M. (2016). Structural basis for leucine sensing by the Sestrin2-mTORC1 pathway. Science 351, 53–58. 10.1126/science.aad2087.

17. Wolfson, R.L., Chantranupong, L., Saxton, R.A., Shen, K., Scaria, S.M., Cantor, J.R., and Sabatini, D.M. (2016). Sestrin2 is a leucine sensor for the mTORC1 pathway. Science 351, 43–48. 10.1126/science.aab2674.

18. Wolfson, R.L., and Sabatini, D.M. (2017). The Dawn of the Age of Amino Acid Sensors for the mTORC1 Pathway. Cell Metab 26, 301–309. 10.1016/j.cmet.2017.07.001.

19. Inglis, A.J., Masson, G.R., Shao, S., Perisic, O., McLaughlin, S.H., Hegde, R.S., and Williams, R.L. (2019). Activation of GCN2 by the ribosomal P-stalk. Proc Natl Acad Sci U S A 116, 4946–4954. 10.1073/pnas.1813352116.

20. Ishimura, R., Nagy, G., Dotu, I., Chuang, J.H., and Ackerman, S.L. (2016). Activation of GCN2 kinase by ribosome stalling links translation elongation with translation initiation. Elife 5. 10.7554/eLife.14295.

21. Masson, G.R. (2019). Towards a model of GCN2 activation. Biochem Soc Trans 47, 1481–1488. 10.1042/BST20190331.

22. Misra, J., Carlson, K.R., Spandau, D.F., and Wek, R.C. (2024). Multiple mechanisms activate GCN2 eIF2 kinase in response to diverse stress conditions. Nucleic Acids Res 52, 1830–1846. 10.1093/nar/gkae006.

23. Misra, J., Holmes, M.J. E. T.M., Langevin, M., Kim, H.G., Carlson, K.R., Watford, M., Dong, X.C., Anthony, T.G., and Wek, R.C. (2021). Discordant regulation of eIF2 kinase GCN2 and mTORC1 during nutrient stress. Nucleic Acids Res 49, 5726–5742. 10.1093/nar/gkab362.

24. Cangelosi, A.L., Puszynska, A.M., Roberts, J.M., Armani, A., Nguyen, T.P., Spinelli, J.B., Kunchok, T., Wang, B., Chan, S.H., Lewis, C.A., et al. (2022). Zonated leucine sensing by Sestrin-mTORC1 in the liver controls the response to dietary leucine. Science 377, 47–56. 10.1126/science.abi9547.

25. Chen, J., Ou, Y., Luo, R., Wang, J., Wang, D., Guan, J., Li, Y., Xia, P., Chen, P.R., and Liu, Y. (2021). SAR1B senses leucine levels to regulate mTORC1 signalling. Nature. 10.1038/s41586-021-03768-w.

26. Kim, J., and Guan, K.L. (2019). mTOR as a central hub of nutrient signalling and cell growth. Nat Cell Biol 21, 63–71. 10.1038/s41556-018-0205-1.

27. Kim, Y., Sundrud, M.S., Zhou, C., Edenius, M., Zocco, D., Powers, K., Zhang, M., Mazitschek, R., Rao, A., Yeo, C.Y., et al. (2020). Aminoacyl-tRNA synthetase inhibition activates a pathway that branches from the canonical amino acid response in mammalian cells. Proc Natl Acad Sci U S A 117, 8900–8911. 10.1073/pnas.1913788117.

28. Lee, M., Kim, J.H., Yoon, I., Lee, C., Fallahi Sichani, M., Kang, J.S., Kang, J., Guo, M., Lee, K.Y., Han, G., et al. (2018). Coordination of the leucine-sensing Rag GTPase cycle by leucyl-tRNA synthetase in the mTORC1 signaling pathway. Proc Natl Acad Sci U S A 115, E5279–E5288. 10.1073/pnas.1801287115.

29. Carroll, B., Maetzel, D., Maddocks, O.D., Otten, G., Ratcliff, M., Smith, G.R., Dunlop, E.A., Passos, J.F., Davies, O.R., Jaenisch, R., et al. (2016). Control of TSC2-Rheb signaling axis by arginine regulates mTORC1 activity. Elife 5. 10.7554/eLife.11058.

30. Cui, Z., Joiner, A.M.N., Jansen, R.M., and Hurley, J.H. (2023). Amino acid sensing and lysosomal signaling complexes. Curr Opin Struct Biol 79, 102544. 10.1016/j.sbi.2023.102544.

31. Fernandes, S.A., Angelidaki, D.D., Nuchel, J., Pan, J., Gollwitzer, P., Elkis, Y., Artoni, F., Wilhelm, S., Kovacevic-Sarmiento, M., and Demetriades, C. (2024). Spatial and functional separation of mTORC1 signalling in response to different amino acid sources. Nat Cell Biol 26, 1918–1933. 10.1038/s41556-024-01523-7.

32. Livneh, I., Cohen-Kaplan, V., Fabre, B., Abramovitch, I., Lulu, C., Nataraj, N.B., Lazar, I., Ziv, T., Yarden, Y., Zohar, Y., et al. (2023). Regulation of nucleo-cytosolic 26S proteasome translocation by aromatic amino acids via mTOR is essential for cell survival under stress. Mol Cell 83, 3333–3346 e3335. 10.1016/j.molcel.2023.08.016.

33. Livneh, I., Fabre, B., Goldhirsh, G., Lulu, C., Zinger, A., Shammai Vainer, Y., Kaduri, M., Dahan, A., Ziv, T., Schroeder, A., et al. (2024). Inhibition of nucleo-cytoplasmic proteasome translocation by the aromatic amino acids or silencing Sestrin3-their sensing mediator-is tumor suppressive. Cell Death Differ 31, 1242–1254. 10.1038/s41418-024-01370-x.

34. Pu, Y., Li, L., Peng, H., Liu, L., Heymann, D., Robert, C., Vallette, F., and Shen, S. (2023). Drug-tolerant persister cells in cancer: the cutting edges and future directions. Nat Rev Clin Oncol 20, 799–813. 10.1038/s41571-023-00815-5.

35. Shorthouse, D., Bradley, J., Critchlow, S.E., Bendtsen, C., and Hall, B.A. (2022). Heterogeneity of the cancer cell line metabolic landscape. Mol Syst Biol 18, e11006. 10.15252/msb.202211006.

36. Liang, X., Zhang, L., Natarajan, S.K., and Becker, D.F. (2013). Proline mechanisms of stress survival. Antioxid Redox Signal 19, 998–1011. 10.1089/ars.2012.5074.

37. Todorova, P.K., Jackson, B.T., Garg, V., Paras, K.I., Brunner, J.S., Bridgeman, A.E., Chen, Y., Baksh, S.C., Yan, J., Hadjantonakis, A.K., and Finley, L.W.S. (2024). Amino acid intake strategies define pluripotent cell states. Nat Metab 6, 127–140. 10.1038/s42255-023-00940-6.

38. Torrence, M.E., MacArthur, M.R., Hosios, A.M., Valvezan, A.J., Asara, J.M., Mitchell, J.R., and Manning, B.D. (2021). The mTORC1-mediated activation of ATF4 promotes protein and glutathione synthesis downstream of growth signals. Elife 10. 10.7554/eLife.63326.

39. Pakos-Zebrucka, K., Koryga, I., Mnich, K., Ljujic, M., Samali, A., and Gorman, A.M. (2016). The integrated stress response. EMBO Rep 17, 1374–1395. 10.15252/embr.201642195.

40. Darnell, A.M., Subramaniam, A.R., and O’Shea, E.K. (2018). Translational Control through Differential Ribosome Pausing during Amino Acid Limitation in Mammalian Cells. Mol Cell 71, 229–243 e211. 10.1016/j.molcel.2018.06.041.

41. Poltorack, C.D., and Dixon, S.J. (2022). Understanding the role of cysteine in ferroptosis: progress & paradoxes. FEBS J 289, 374–385. 10.1111/febs.15842.

42. Zheng 郑嘉烁, J., and Conrad, M. (2024). Ferroptosis: when metabolism meets cell death. Physiol Rev. 10.1152/physrev.00031.2024.

43. Wang, J., Alexander, P., Wu, L., Hammer, R., Cleaver, O., and McKnight, S.L. (2009). Dependence of mouse embryonic stem cells on threonine catabolism. Science 325, 435–439. 10.1126/science.1173288.

44. Tarangelo, A., Rodencal, J., Kim, J.T., Magtanong, L., Long, J.Z., and Dixon, S.J. (2022). Nucleotide biosynthesis links glutathione metabolism to ferroptosis sensitivity. Life Sci Alliance 5. 10.26508/lsa.202101157.

45. Bohm, R., Imseng, S., Jakob, R.P., Hall, M.N., Maier, T., and Hiller, S. (2021). The dynamic mechanism of 4E-BP1 recognition and phosphorylation by mTORC1. Mol Cell. 10.1016/j.molcel.2021.03.031.

46. Kang, S.A., Pacold, M.E., Cervantes, C.L., Lim, D., Lou, H.J., Ottina, K., Gray, N.S., Turk, B.E., Yaffe, M.B., and Sabatini, D.M. (2013). mTORC1 phosphorylation sites encode their sensitivity to starvation and rapamycin. Science 341, 1236566. 10.1126/science.1236566.

47. Shimobayashi, M., and Hall, M.N. (2014). Making new contacts: the mTOR network in metabolism and signalling crosstalk. Nat Rev Mol Cell Biol 15, 155–162. 10.1038/nrm3757.

48. Thoreen, C.C., Kang, S.A., Chang, J.W., Liu, Q., Zhang, J., Gao, Y., Reichling, L.J., Sim, T., Sabatini, D.M., and Gray, N.S. (2009). An ATP-competitive mammalian target of rapamycin inhibitor reveals rapamycin-resistant functions of mTORC1. J Biol Chem 284, 8023–8032. 10.1074/jbc.M900301200.

49. Oren, Y., Tsabar, M., Cuoco, M.S., Amir-Zilberstein, L., Cabanos, H.F., Hutter, J.C., Hu, B., Thakore, P.I., Tabaka, M., Fulco, C.P., et al. (2021). Cycling cancer persister cells arise from lineages with distinct programs. Nature 596, 576–582. 10.1038/s41586-021-03796-6.

50. Hangauer, M.J., Viswanathan, V.S., Ryan, M.J., Bole, D., Eaton, J.K., Matov, A., Galeas, J., Dhruv, H.D., Berens, M.E., Schreiber, S.L., et al. (2017). Drug-tolerant persister cancer cells are vulnerable to GPX4 inhibition. Nature 551, 247–250. 10.1038/nature24297.

51. Takahashi, N., Cho, P., Selfors, L.M., Kuiken, H.J., Kaul, R., Fujiwara, T., Harris, I.S., Zhang, T., Gygi, S.P., and Brugge, J.S. (2020). 3D Culture Models with CRISPR Screens Reveal Hyperactive NRF2 as a Prerequisite for Spheroid Formation via Regulation of Proliferation and Ferroptosis. Mol Cell 80, 828–844 e826. 10.1016/j.molcel.2020.10.010.

52. Sullivan, M.R., Danai, L.V., Lewis, C.A., Chan, S.H., Gui, D.Y., Kunchok, T., Dennstedt, E.A., Vander Heiden, M.G., and Muir, A. (2019). Quantification of microenvironmental metabolites in murine cancers reveals determinants of tumor nutrient availability. Elife 8. 10.7554/eLife.44235.

53. Saxton, R.A., and Sabatini, D.M. (2017). mTOR Signaling in Growth, Metabolism, and Disease. Cell 168, 960–976. 10.1016/j.cell.2017.02.004.

54. Vettore, L., Westbrook, R.L., and Tennant, D.A. (2020). New aspects of amino acid metabolism in cancer. Br J Cancer 122, 150–156. 10.1038/s41416-019-0620-5.

55. Bar-Peled, L., Chantranupong, L., Cherniack, A.D., Chen, W.W., Ottina, K.A., Grabiner, B.C., Spear, E.D., Carter, S.L., Meyerson, M., and Sabatini, D.M. (2013). A Tumor suppressor complex with GAP activity for the Rag GTPases that signal amino acid sufficiency to mTORC1. Science 340, 1100–1106. 10.1126/science.1232044.

56. Howden, A.J.M., Hukelmann, J.L., Brenes, A., Spinelli, L., Sinclair, L.V., Lamond, A.I., and Cantrell, D.A. (2019). Quantitative analysis of T cell proteomes and environmental sensors during T cell differentiation. Nat Immunol 20, 1542–1554. 10.1038/s41590-019-0495-x.

57. Hukelmann, J.L., Anderson, K.E., Sinclair, L.V., Grzes, K.M., Murillo, A.B., Hawkins, P.T., Stephens, L.R., Lamond, A.I., and Cantrell, D.A. (2016). The cytotoxic T cell proteome and its shaping by the kinase mTOR. Nat Immunol 17, 104–112. 10.1038/ni.3314.

58. Marchingo, J.M., Sinclair, L.V., Howden, A.J., and Cantrell, D.A. (2020). Quantitative analysis of how Myc controls T cell proteomes and metabolic pathways during T cell activation. Elife 9. 10.7554/eLife.53725.

59. Van de Velde, L.A., and Murray, P.J. (2016). Proliferating Helper T Cells Require Rictor/mTORC2 Complex to Integrate Signals from Limiting Environmental Amino Acids. J Biol Chem 291, 25815–25822. 10.1074/jbc.C116.763623.

60. Van de Velde, L.A., Subramanian, C., Smith, A.M., Barron, L., Qualls, J.E., Neale, G., Alfonso-Pecchio, A., Jackowski, S., Rock, C.O., Wynn, T.A., and Murray, P.J. (2017). T Cells Encountering Myeloid Cells Programmed for Amino Acid-dependent Immunosuppression Use Rictor/mTORC2 Protein for Proliferative Checkpoint Decisions. J Biol Chem 292, 15–30. 10.1074/jbc.M116.766238.

61. Conlon, M., Poltorack, C.D., Forcina, G.C., Armenta, D.A., Mallais, M., Perez, M.A., Wells, A., Kahanu, A., Magtanong, L., Watts, J.L., et al. (2021). A compendium of kinetic modulatory profiles identifies ferroptosis regulators. Nat Chem Biol 17, 665–674. 10.1038/s41589-021-00751-4.

62. Murray, P.J. (2016). Amino acid auxotrophy as a system of immunological control nodes. Nat Immunol 17, 132–139. 10.1038/ni.3323.

63. Zeitler, L., and Murray, P.J. (2023). IL4i1 and IDO1: Oxidases that control a tryptophan metabolic nexus in cancer. J Biol Chem 299, 104827. 10.1016/j.jbc.2023.104827.

64. An, H., Ordureau, A., Korner, M., Paulo, J.A., and Harper, J.W. (2020). Systematic quantitative analysis of ribosome inventory during nutrient stress. Nature 583, 303–309. 10.1038/s41586-020-2446-y.

65. Klann, K., Tascher, G., and Munch, C. (2020). Functional Translatome Proteomics Reveal Converging and Dose-Dependent Regulation by mTORC1 and eIF2alpha. Mol Cell 77, 913–925 e914. 10.1016/j.molcel.2019.11.010.

66. Bruggenthies, J.B., Fiore, A., Russier, M., Bitsina, C., Brotzmann, J., Kordes, S., Menninger, S., Wolf, A., Conti, E., Eickhoff, J.E., and Murray, P.J. (2022). A cell-based chemical-genetic screen for amino acid stress response inhibitors reveals torins reverse stress kinase GCN2 signaling. J Biol Chem, 102629. 10.1016/j.jbc.2022.102629.

67. Schmidt, E.K., Clavarino, G., Ceppi, M., and Pierre, P. (2009). SUnSET, a nonradioactive method to monitor protein synthesis. Nat Methods 6, 275–277. 10.1038/nmeth.1314.

68. Dobin, A., Davis, C.A., Schlesinger, F., Drenkow, J., Zaleski, C., Jha, S., Batut, P., Chaisson, M., and Gingeras, T.R. (2013). STAR: ultrafast universal RNA-seq aligner. Bioinformatics 29, 15–21. 10.1093/bioinformatics/bts635.

69. Anders, S., Pyl, P.T., and Huber, W. (2015). HTSeq--a Python framework to work with high-throughput sequencing data. Bioinformatics 31, 166–169. 10.1093/bioinformatics/btu638.

70. Anders, S., and Huber, W. (2010). Differential expression analysis for sequence count data. Genome Biology 11, R106. 10.1186/gb-2010-11-10-r106.

71. Humphrey, S.J., Karayel, O., James, D.E., and Mann, M. (2018). High-throughput and high-sensitivity phosphoproteomics with the EasyPhos platform. Nat Protoc 13, 1897–1916. 10.1038/s41596-018-0014-9.

72. Tyanova, S., Temu, T., Sinitcyn, P., Carlson, A., Hein, M.Y., Geiger, T., Mann, M., and Cox, J. (2016). The Perseus computational platform for comprehensive analysis of (prote)omics data. Nat Methods 13, 731–740. 10.1038/nmeth.3901.

73. Bekker-Jensen, D.B., Bernhardt, O.M., Hogrebe, A., Martinez-Val, A., Verbeke, L., Gandhi, T., Kelstrup, C.D., Reiter, L., and Olsen, J.V. (2020). Rapid and site-specific deep phosphoproteome profiling by data-independent acquisition without the need for spectral libraries. Nat Commun 11, 787. 10.1038/s41467-020-14609-1.

